# Subunit positioning and diversity of the nematode levamisole-sensitive acetylcholine receptor

**DOI:** 10.1101/2024.05.05.592571

**Authors:** JD Noonan, TB Duguet, CJ Handy-Hart, RN Beech

## Abstract

The helminth levamisole-sensitive acetylcholine receptor (L-AChR) is a historically significant drug target. This heteromeric receptor belongs to the larger class of pentameric ligand-gated ion channels that has recently expanded within the nematodes and therefore provides the opportunity to study specific examples of how worm receptors evolve. The *Caenorhabditis elegans* L-AChR can exist in two forms. The first contains three alpha subunits: ACR-13, UNC-38, and UNC-63, and two non-alpha subunits UNC-29 and LEV-1. The second can be formed by replacing ACR-13 with its paralog ACR-8, from either *C. elegans* or the closely related *Haemonchus contortus.* This form no longer requires LEV-1, demonstrating an evolved functional difference between ACR-8 and ACR-13. Since the properties of a channel depend on its composition and organization, knowing the underlying mechanisms regulating this would provide valuable insight into one of the least understood aspects of this important drug target. The goal of this study was to identify the subunit stoichiometry of the L-AChR and further elucidate the functional divergence of alpha subunits ACR-8 and ACR-13. Using a series of subunit concatemers, we determined the arrangement of three *C. elegans* L-AChR subunits in the following N- to C-terminal order; UNC-38 – LEV-1 – UNC-63. We were unable to order the ACR-13 and UNC-29 subunits unambiguously. Additionally, our concatemers provide support for the sequential model of subunit assembly whereby a trimer is formed first, consisting of UNC-38 – LEV-1 – UNC-63, followed by individual addition of the remaining two subunits ACR-13 and UNC-29. Using *C. elegans* and *H. contortus* subunit admixtures, we show that replacing ACR-13 with ACR-8 alleviates the requirement for five distinct subunits, as a functional receptor can be measured in the absence of non-alpha LEV-1. We confirm that UNC-29 can replace LEV-1 and that the intracellular loop determines this positional plasticity. This work provides the first evidence of the quaternary structure of the L-AChR which is necessary for future studies on this drug target. It also confirms that subunit arrangement is determined first during receptor heteromerization, followed by functional fine-tuning once subunit types have defined positions.

## 1. Introduction

The levamisole-sensitive acetylcholine receptor (L-AChR) of the free-living model nematode *Caenorhadbitis elegans* has been the focus of research into the development of the nervous system for the past 40 years (Lewis et al., 1980b, 1980a; Jospin et al., 2022). This receptor belongs to a larger protein superfamily known as the pentameric ligand-gated ion channels (pLGICs). The pLGICs are the targets of many anthelmintics and so a detailed understanding of this receptor family is of pharmacological significance (Wolstenholme, 2011; Martin et al., 2012). pLGICs contain five subunits arranged in a radially symmetrical complex with a central ion-permeable pore (Dacosta and Baenziger, 2013). Each subunit is structurally similar and consists of a large extracellular domain, where ligand binding occurs, followed by four transmembrane domains that form the pore, and a large intracellular loop (Dacosta and Baenziger, 2013; Wu et al., 2015).

The ligand-binding and gating mechanisms of pLGICs are highly conserved. Ligands bind at the interface between two subunits, with each subunit contributing one face of the pocket (Corringer et al., 2000; Pless and Sivilotti, 2018). The principle face contains a vicinal cysteine motif that is central in gating, the complementary face does not contain this amino acid signature (Corringer et al., 2000; Dacosta and Baenziger, 2013). In nematodes, alpha subunits contain the vicinal cysteine motif, whereas non-alpha subunits do not (Beech and Neveu, 2015). Homomeric channels are therefore composed of alpha subunits, whereas heteromeric receptors are composed of alpha and non-alpha subunits. In a heteromeric receptor the alpha subunit provides the principle face of the binding pocket and the non-alpha subunit provides the complementary face (Corringer et al., 2000; Jones and Sattelle, 2003). Despite there being five subunit interfaces in a receptor, two ligand molecules bind in both homomeric and heteromeric receptors and this asymmetry provides instability required for gating (Dacosta and Baenziger, 2013; Mowrey et al., 2013). Solved structures of cationic heteropentameric receptors reveal subunit stoichiometries with this asymmetry (as examples: PDB 2BG9 & 5KXI) (Unwin, 2005; Morales-Perez et al., 2016).The position that a subunit occupies in a heteromeric receptor is therefore critical for receptor function and must be strictly maintained.

A phylogeny of L-AChR subunits shows that the division of alpha subunits (*acr-8, acr-13, unc-38* and *unc-63*) from non-alpha subunits (*lev-1* and *unc-29*) occurred before the duplication and divergence producing the different specialized subunits (Jones et al., 2007; Jones and Sattelle, 2008; Beech and Neveu, 2015). Within the alpha subunits, paralogous pairs *acr-8* and *acr-13,* as well as *unc-63* and *unc-38* derive from duplications. The *C. elegans* L-AChR (Cel-L-AChR) can exist in multiple forms, the first (Cel-L-AChR-1) contains three alpha subunits ACR-13, UNC-38, and UNC-63, and two non-alpha subunits UNC-29 and LEV-1 (Boulin et al., 2008). Removal of any one subunit leads to loss of the receptor (Boulin et al., 2008), emphasizing the defined exclusive positioning of each subunit. This is an extreme example where five different genes encode the receptor. The other Cel-L-AChRs can be formed by replacing ACR-13 with its paralog ACR-8. In this case two receptor types can be formed; one with LEV-1 (Cel-L-AChR-2.1) and one without LEV-1 (Cel-L-AChR-2.2) (Blanchard et al., 2018). This demonstrates an evolved functional difference between ACR-8 and ACR-13; ACR-13 requires LEV-1 whereas ACR-8 does not. Despite over 40 years of research into the *C. elegans* L-AChR subunits (Lewis et al., 1980a, 1980b), the relative subunit positioning within the receptor is still not known.

The L-AChRs identified from parasitic nematodes exhibit more flexible subunit arrangements than observed in*C. elegans*. Two different *Ascaris suum* L-AChRs can be produced from only UNC-38 and UNC-29 (Williamson et al., 2009). In *Oesophagostomum dentatum,* four different L-AChRs can be made from ACR-8, UNC-38, UNC-63 and UNC-29 (Buxton et al., 2014). Meanwhile, *Haemonchus contortus* has four full length copies of the non-alpha subunit *unc-29* (*unc-29.1 - unc-29.4*) and at least four different L-AChRs can be produced. In *H. contortus*, UNC-29.1 produces a functional L-AChR without LEV-1, whereas UNC-29.2 does not (Boulin et al., 2011; Duguet et al., 2016). In all these cases where fewer than five different subunits are present, at least one of the subunits must be present twice, occupying more than one position, a striking difference compared to the Cel-L-AChR-1. Therefore, diversity in receptor composition and subunit positioning vary between closely related species.

Given this receptor complexity, it is easy to speculate that subunits in heteromeric receptors evolve to specialize their roles, leading to co-adaptation between the subunits restricted to a specific arrangement and compatibility. The L-AChR of nematodes provides an experimentally accessible system to investigate this; these are examples where non-alpha subunits (UNC-29 paralogs) differ in their requirement for another non-alpha subunit (LEV-1) and alpha subunits (ACR-8 and ACR-13) also differ in their requirement for the same non-alpha subunit (LEV-1) (Duguet et al., 2016). Combining subunits from different closely related species into admixture receptor combinations would disrupt previously evolved inter-subunit interactions and provide a means to investigate the mechanism. However, interpreting the results of cross-species admixtures is limited because the L-AChR subunit stoichiometry remains unknown. Identifying the arrangement of the subunits would thus provide crucial insight needed for understanding previous and current L-AChR studies (Boulin et al., 2008, 2011; Williamson et al., 2009; Buxton et al., 2014; Duguet et al., 2016; Blanchard et al., 2018). The goal of this study was to therefore examine subunit stoichiometry of the Cel-L-AChR and further elucidate the functional divergence of alpha subunits ACR-8 and ACR-13 and non-alpha UNC-29 paralogs.

## 2. Methods

### 2.1 Subunit cloning

Subunit nomenclature used here follows established nematode ion channel subunit naming, subunits in italics refer to subunit genes and capitalized subunits refer to proteins (Beech et al., 2010). The expression clones for AChR subunits and ancillary proteins from *H. contortus* were available in the pTB207 vector that contains the beta-globin 3’ UTR, while expression clones for *C. elegans* (except for *Cel-acr-8*) were kindly provided by C. Neveu (Boulin et al., 2008).

The cDNA clones employed in this research correspond to the following sequences: Cel-lev-1 (NP_502534), Cel-acr-13 (formerly lev-8) (NP_509932), Cel-unc-29 (NP_492399), Cel-unc-38 (NP_491472), and Cel-unc-63 (NP_491533). Additionally, they include Hco-unc-29.1 (GU060980), Hco-unc-29.2 (GU060981), Hco-unc-29.3 (GU060982), Hco-unc-29.4 (GU060983), Hco-unc-38 (GU060984), Hco-unc-63a (GU060985), Hco-acr-8 (EU006785) and the chaperon sequences Hco-unc-50 (HQ116822), Hco-unc-74 (HQ116821), and Hco-ric-3.1 (HQ116823). All newly cloned sequences were inserted into the pTD2 expression vector consisting of the pTB207 plasmid supplemented with the *Xenopus laevis* beta-globin 5’ UTR (Duguet et al., 2016).

#### 2.1.1 Cel-acr-8

*C. elegans acr-8* was cloned from worm cDNA using primers 5’- GCGGCCGCAACCGACAATACCATGAACTTCGC -3’ and 5’- GGGCCCACATATTCATTGATATTTGATGGG –3’. The forward primer introduces a *NotI* site at the N-terminal region while the reverse primer introduces an *ApaI* site after the stop codon. The PCR product was purified through an 0.8% agarose gel, recovered (ZymoResearch, USA) and ligated into pGEM-T (Promega, USA). The ligation reaction was transformed into DH5α *Escherichia coli* (ThermoFisher Scientific, USA) grown on LB plates with 100 μM Ampicilin (ThermoFisher Scientific, USA). Plasmids were purified (BioBasic, USA) and sent for sequencing (Molecular Cloning Lab, USA). The correct *acr-8* clone was then transferred to the pTD2 expression vector. It was first digested with *NotI* and *ApaI* (Fast Digest, ThermoFisher Scientific, USA) and ligated into the pTD2 plasmid. The ligation mixture was then transformed into DH5α *Escherichia coli* (ThermoFisher Scientific, USA) grown on LB plates with 100 μM Ampicilin (ThermoFisher Scientific, USA).

#### 2.1.2 Concatemers

The *C. elegans* L-AChR subunit cDNA sequences were re-amplified by PCR with primers containing restriction sites (Supplemental Figure 1). This allowed precise positioning of subunits into the pTB207 plasmid using an appropriate mix of FastDigest (ThermoFisher Scientific, USA) restriction enzymes. For generating dimers, the subunits in the first position were amplified with *Not*I (5’) and *Xba*I (3’). The *Xba*I-containing reverse primer was designed to eliminate the stop codon for continuous translation from the first subunit to the second subunit. The subunit sequence in the second position was amplified with *Xba*I and *Apa*I restriction sites at the 5’ and 3’ ends, respectively, and the *Xba*I-containing primer was designed to remove the signal peptide. To create trimers, the second-position subunits were amplified to remove the signal peptide and stop codon, introducing *Xba*I (5’) and *Afl*II (3’) restriction sites. The third subunit sequence, lacking a signal peptide but containing a stop codon, was amplified with primers incorporating *Afl*II (5’) and *Apa*I (3’) restriction sites. As only two tetrameric structures were generated, the [*unc-38:lev-1:unc-63]* trimer structure served as a template. The trimer was digested with *Afl*II and *Apa*I to remove the third subunit sequence. The desired third subunit was then amplified (without signal peptide/stop codon) and flanked with *Afl*II and *Avr*II restriction sites at the 5’ and 3’ ends, respectively. The fourth subunit was cloned without a signal peptide and supplemented with the *Avr*II (5’) and *Apa*I (3’) restriction sites. Using a similar cloning strategy, pentameric constructs were cloned from the [*unc-38:lev-1:unc-63*] trimeric template. The fourth subunit was amplified without a signal peptide sequence and stop codon, flanked with *Avr*II (5’) and *Xba*I (3’) restriction sites. The fifth subunit sequence, depleted of its signal peptide sequence, was amplified with *Xba*I and *Apa*I-containing primers.

Models of the dimer concatemers were made to ensure they did not alter the subunit structure. These were based on subunits A and B of the human α4β2 acetylcholine receptor (PDB: 5KXI) (Morales-Perez et al., 2016). The alignment was constructed by hand, based on the conserved secondary and tertiary structural elements. The models were developed using the hm_build macro of YASARA 23.12.24 (Krieger et al., 2002). In each case, the C-terminal of the principle face subunit was covalently attached to the N-terminal of the complementary subunit face (Supplementary Figure 2).

#### 2.1.3 Chimeras

Chimeras in the pTD2 expression vector were purchased from Gene Universal Inc. (Newark, USA). Sequences were designed by exchanging the intracellular loop between *Hco-unc-29.1* and *Hco-unc-29.2*. Chim-1 refers to the *unc-29.1* subunit sequence containing the *unc-29.2* intracellular loop and Chim-2 refers to *unc-29.2* subunit sequence containing the *unc-29.1* intracellular loop.

### 2.2 cRNA preparation

cRNA was prepared by either linearizing the plasmid DNA with *NheI* or with a PCR using forward primer pTD2F (5’-TTGGCACCAAAATCAACGGG – 3’) and reverse primer SP6 (5’- ATTTAGGTGACACTATAG -3’) (SuperFi DNA Polymerase, ThermoFischer Scientific, USA). In both cases the DNA template used for the cRNA reaction contained the gene of interest the poly-A tail, the T7 promoter sequence and the *X. leavis* beta-globin UTRs. *In vitro* transcription with the mMESSAGE mMACHINE T7 kit (Ambion, USA) was carried out following manufacturers protocol. Precipitated cRNA was dissolved in RNase-free water and its concentration was measured with a Nanodrop spectrophotometer (ThermoFisher Scientific, USA).

### 2.3 Frog oocyte extraction & injection

*X. laevis* were used in this study in accordance with the McGill Animal Use Protocol AUP-2015-7758 authorized by the McGill University Animal Care Committee of the Office of Research Ethics and Compliance. Oocyte extraction surgeries were carried out by trained personnel following standard protocol. Adult female frogs were anaesthetized in 0.15% tricaine mesylate (MS-222, Sigma-Aldrich, USA) (pH 7.3 with NaHCO3). A <1 cm incision was made on the side of the abdomen and oocytes extracted using fine tweezers. The incision was sutured, and the frog returned to the Xenoplus Housing System (Technoplast, Italy). Frogs were monitored daily for two weeks post-surgery and returned to the colony tank. Extracted oocytes were prepared according to standard protocol (Goldin, 1991). The lobes were placed in Ca^2+^-free OR2 solution (82 mM NaCl, 2 mM KCl, 1 mM MgCl2, 5 mM HEPES, pH 7.3) and divided into clusters of <10 oocytes using tweezers. Oocytes were then incubated in 10 mg/mL collagenase type II (Sigma-Aldrich, USA) for 90 minutes, washed in Ca^2+^-free OR2, placed in ND96 solution (NaCl 96 mM, KCl 2 mM, CaCl_2_ 1.8 mM, MgCl_2_ 1 mM and HEPES 5 mM, pH 7.3) supplemented with sodium pyruvate 2.5 mM (Forrester et al., 2003) and incubated at 18°C until cRNA injection.

Oocytes were injected with 50 nL (12.5 ng per injected gene) of the desired cRNA subunit and accessory protein combination. All combinations were co-injected with the *H. contortus* accessory proteins *ric-3.1, unc-50* and *unc-74* (Halevi et al., 2002; Eimer et al., 2007; Boulin et al., 2008). Oocytes were immersed in ND96 solution and injected via Nanoject (Drummond Scientific, USA) using pulled glass capillaries (World Precision Instruments, USA). Following injection oocytes were incubated in ND96 buffer at 18°C for 4-5 days until electrophysiology.

### 2.4 Two-electrode voltage clamp electrophysiology

Acetylcholine (ACh) (Sigma-aldrich, USA) and levamisole ((-)-tetramisole) hydrochloride (LEV) (Sigma-aldrich, USA) ligand solutions were dissolved in Ringers solution (NaCl 100 mM, KCl 2.5 mM, CaCl2 1 mM, HEPES 5 mM, pH 7.3) to a final recording concentration of 100 μM. Oocytes were placed in an RC3Z bath chamber (Harvard Apparatus, USA) submerged in Ringers solution. The bath chamber was grounded with a 3M KCl agar bridge. A perfusion system containing Ringer solution and both ligand solutions was connected to the chamber allowing solutions to perfuse over the oocyte at an approximate rate of 1.2 m/s. Pulled glass capillaries (World Precision Instruments, USA) were filled with 3M KCl and the tips clipped with tweezers to a resistance between 0.5 and 5 MΩ. Measurements were made using an Axon Axoclamp 900A amplifier and Digidata 1550M. Oocytes were voltage-clamped at -60 mV and first exposed to Ringers recording solution and current allowed to stabilize, those that did not stabilize were discarded. Once stabilized, ligand solutions were perfused over the oocytes and current responses were measured until a current peak was measured followed by a wash with the Ringer solution. When the membrane current returned to baseline, the next ligand was applied, and peak current again recorded. Data was analyzed using Clampex 9.2 (Axon Instruments, USA) and statistical analysis and graphs produced in Prism (GraphPad version 9.1). Unpaired students T-tests were used to compare responses with error bars representing standard error. Representative current traces of oocyte recordings were filtered in Clampfit using the lowband pass at 40 Hz cutoff.

## 3. Results

### 3.1 Quaternary structure of the Cel-L-AChR

A defined subunit arrangement of the L-AChR is necessary to interpret admixtures presented here and elsewhere (Boulin et al., 2008, 2011; Williamson et al., 2009; Buxton et al., 2014; Duguet et al., 2016; Blanchard et al., 2018). Since more than one type of Cel-L-AChR exists, we sought to determine the subunit arrangement of the receptor with ACR-13. The requirement for this receptor to contain five different subunits meant that each one has an absolute defined position with no uncertainty of any being present more than once.

The approach used here was to concatenate subunits together into one continuous peptide. The C-terminal tail of the first subunit was linked to the N-terminal domain of the next subunit, this was made possible because both are in the extracellular region of the receptor (Dacosta and Baenziger, 2013). Only the first subunit’s signal peptide and the last subunit’s stop codon were retained, making a single open reading frame and restricting the subunits to one defined configuration. These constructs are henceforth referred to as “concatemers” and are denoted by subunits within square brackets separated by colons (ex: [ACR-13:UNC-29]). Given the complexity of producing any pentameric combination from five different subunits, a stepwise approach was used. First, dimer concatemers were made by fusing two subunits together. If the fused subunits are indeed adjacent in the mature receptor then this dimer, along with the three other subunits as individual monomers, would be in the correct configuration and produce a measurable response. The dimers should identify “pairs” of subunits within the complex and fusion of other subunits to the functional dimers, producing trimers, would then evaluate the next subunit in the structure, and so on (Saedi et al., 1991; Green and Claudio, 1993; Kreienkamp et al., 1995; Green and Wanamaker, 1998).

Homology models of the dimeric concatemers based on the human α4β2 nicotinic receptor (PDB: 5KXI) (Morales-Perez et al., 2016) were made to ensure that the fusion did not distort the normal subunit structure (Supplemental Figure 2). These models confirmed that the dimers maintained the normal principle/complementary organization without allowing another free subunit to sit between them. Only UNC-63, had a C-terminal tail of UNC-63 was too short and required the addition of five Ser-Ala-Thr repeats in order to link it to the next subunit. These were chosen as small, non-branched, hydrophylic residues. This linker had no effect on the function of UNC-63 in response to 100 μM ACh (689.9 ± 221.6 nA, n = 5, *p* = 0.194) and 100 μM LEV (372.9 ± 189.1 nA, n = 5, *p* = 5156) (Figure 1F).

**Figure 1.**
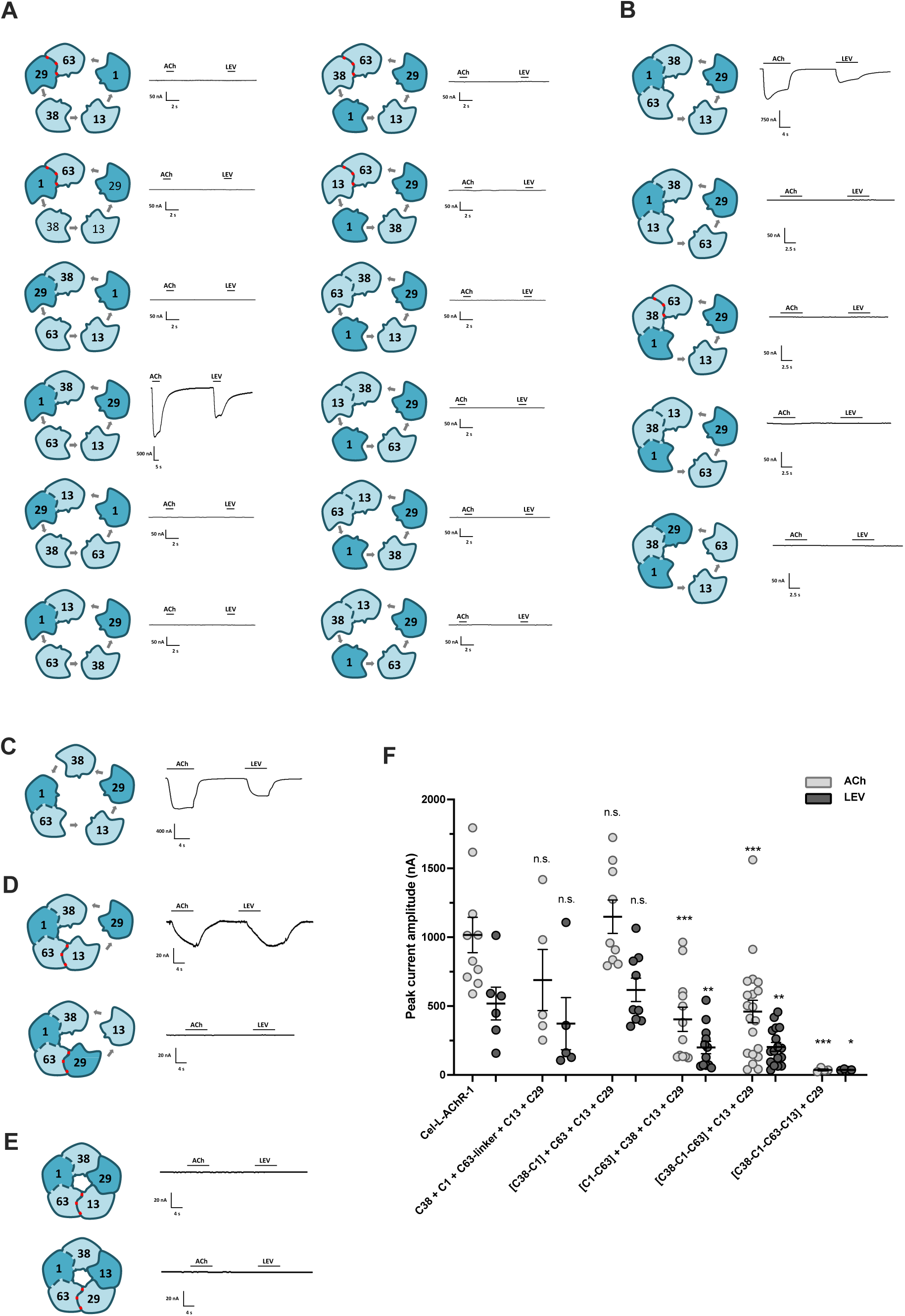
Subunit arrangement of the *C. elegans* levamisole-sensitive acetylcholine receptor. *C. elegans* L-AChR subunits were made into a series of concatemers and assayed for function in oocytes to identify the subunit arrangement of the receptor. (A) Sample recordings of the dimer concatemers show only the [C38-C1] dimer produced current response. (B) Sample recordings of the trimer concatemers that were made based on the success of the [C38-C1] dimer. Only the [C38-C1-C63] trimer condition produced current response. (C) The successful measurement of the [C38-C1-C63] trimer suggested that the [1-63] dimer concatemer not tested previously would also function. The [C1-C63] dimer indeed produced measurable response, supporting the sequential model of subunit assembly. (D) Sample recordings of the tetramer concatemers that were made based on the success of the [C38-C1-C63] trimer. Only the [C38-C1-C63-C13] tetramer produced current response. (E) Sample recordings of the pentamer concatemers that were made based on the [C38-C1-C63] trimer. Neither pentamer concatemer produced current responses. (F) Summary of peak current amplitudes of concatemer + monomer combinations that were functional in oocytes. “C” corresponds to the abbreviation of *C. elegans* followed by the subunit gene number indicated. Brackets represent subunit concatamers. Peak current responses for L-AChR-1 (with ACR13) were 1017.0 ± 128.6 nA, n=10 and 518.8 ± 118.9 nA, n=6; C38 + C1 + C63-linker + C13 + C29 were 689.9 ± 221.6 nA, n=6 and 372.9 ± 189.1 nA, n=5; [C38-C1] + C63 + C13 + C29 were 1149.0 ± 121.3 nA, n=9 and 618.3 ± 84.7 nA, n=9; [C1-C63] + C38+ C13 + C29 were 404.4 ± 88.3 nA, n=12 and 199.6 ± 45.1 nA, n=12; [C38-C1-C63] + C13 + C29 were to 460.5 ± 81.4 nA, n=20 and 204.2 ± 32.0 nA, n=17; [C38-C1-C63-C13] + C29 were 36.9 ± 6.7 nA, n=4 and 37.3 ± 2.5 nA, n=4; for ACh and LEV, respectively. Responses were compared to the native *C. elegans* L-AChR-1 (with ACR-13). n.s.: non-significantly different according to an unpaired Student’s t-test with a P-value p<0.05. * p< 0.05; **p<0.005; ***p<0.0002, **** p<0.0001. Error bars indicate SE. Peak current amplitudes in response to 100 μM ACh (light grey) and 100 μM LEV (dark grey). Concatenated subunits are represented in form of two subunits joined together. The number corresponds to the subunit gene name number, example UNC-38 is the subunit with the number 38. The C refers to *C. elegans*. *C. elegans* non-alpha subunits shown as dark blue and *C. elegans* alpha subunits shown as light blue. Dashed line represents the limit between the two subunits. The red dashed line corresponds to the artificial linker that was added to the C-terminal of Cel-UNC-63. Monomers are showed as individual subunits. See text and Supplemental Figure 1 for concatemer engineering methods. Electrophysiology traces are representative of a single oocyte recording with horizontal bars showing the time period of agonist application.

#### 3.1.1 Dimer concatemers

The total number of possible dimer combinations from 5 different subunits is large, to reduce the number of combinations tested, only twelve, where the first subunit was an alpha subunit, were evaluated initially. Using this approach, at least three dimers should contain subunits in the correct relative position regardless of the real organization of the receptor. Surprisingly, only the [UNC-38:LEV-1] dimer, responded to ACh (1148.8 ± 121.3 nA, n = 9) and LEV (618.3 ± 84.7 nA, n = 9) (Figure 1A). This indicates that UNC-38:LEV-1 is in the correct configuration and one of the first steps of receptor assembly. The other dimers that had subunits in the correct order but failed to produce functional channels may have to join the assembling complex as single subunits (Gu et al., 1991; Green and Wanamaker, 1997).

#### 3.1.2 Trimer concatemers

To identify the next subunit in the structure, a series of trimeric constructs were made based on the [UNC-38:LEV-1] dimer. The trimers concatenated UNC-63 or ACR-13 before UNC-38 or after LEV-1. UNC-29 was only concatenated before UNC-38 since it would be unlikely that both non-alpha subunits would be adjacent to each other. Only the [UNC-38:LEV-1:UNC-63] trimer produced a response to ACh (460.5 ± 81.4, nA n = 20) and LEV (204.2 ± 32.02 nA, n = 17) (Figure 1B).

The results were so far consistent with the sequential model of subunit assembly that states that three subunits initially form a trimer followed by the sequential addition of the remaining two subunits (Green and Claudio, 1993). Based on the success of the [UNC-38:LEV-1:UNC-63] trimer, this model predicts that a [LEV-1:UNC-63] dimer would also produce a functional channel. This combination had not been tested in the initial dimers. A [LEV-1:UNC-63] dimer was subsequently made and indeed produced measurable currents to both ACh (404.4 ± 88.3 nA, n = 12) and LEV (199.6 ± 45.13 nA, n = 12) (Figure 1C), further supporting the sequential model of subunit assembly. However, these responses were significantly lower compared to the [UNC-38:LEV-1] dimer which might suggest a preference for an initial UNC-38:LEV-1 dimer (p<0.0002 for ACh and p<0.001 for LEV).

#### 3.1.3 Tetramer and pentamer concatemers

Success of the [UNC-38:LEV-1:UNC-63] trimer meant only two possibilities were left to determine; either UNC-29 followed by ACR-13 or *vice versa*. Therefore, two tetramer concatemers were made with either UNC-29 or ACR-13 in the fourth position with the fifth subunit as a monomer. The [UNC-38:LEV-1:UNC-63:UNC-29] tetramer produced no response to ACh or LEV (n=5) (Figure 1D) while the [UNC-38:LEV-1:UNC-63:ACR-13] tetramer produced small current responses to ACh (36.96 ± 6.65 nA, n = 4) and LEV (37.28 ± 2.46 nA, n = 4) (Figure 1D). These ACh responses were a loss of 96.5 ± 0.6 % of the ACh-evoked currents measured from the native Cel-L-AChR-1 and demonstrated a significant loss of signal as the concatemer chain length increases. Two pentamer concatemers were made to further test both arrangements of ACR-13 and UNC-29. Unfortunately, neither [UNC-38:LEV-1:UNC-63:ACR-13:UNC-29] nor [UNC-38:LEV-1:UNC-63:UNC-29:ACR-13] produced detectable responses (Figure 1E). A summary of the functional concatemers is shown in Figure 1F. Based on these concatemers we propose the subunit arrangement of the Cel-L-AChR-1 as from N terminal to C terminal: UNC-38:LEV-1:UNC-63:ACR-13:UNC-29. We claim a demonstration of the first of these subunits, but the complete pentamer would require further confirmation.

### 3.2 The non-alpha UNC-29 subunit replaces LEV-1

When ACR-13 is replaced by ACR-8 in the Cel-L-AChR, the receptor no longer requires LEV-1 (Blanchard et al., 2018). Since five subunits are needed for the pentameric structure, one of the other subunits must be present twice. Subunit phylogeny would predict that the other non-alpha subunit, UNC-29, would replace LEV-1 (Beech and Neveu, 2015), however this has not been confirmed. We sought to examine this using the concatemer dimers described.

If UNC-29 can replace LEV-1, then a dimer of [UNC-38:UNC-29] should produce a functional receptor in combination with ACR-8, but not ACR-13 when LEV-1 is omitted. During these recordings the control conditions of the dimer concatemers [UNC-38:LEV-1] with monomers UNC- 63 + UNC-29 + ACR-13 or UNC-63 + UNC-29 + ACR-8 all produced small current responses (∼200 - 400 nA) (Figure 2A - C). Considering the [UNC-38:LEV-1] + UNC-63 + UNC-29 + ACR- 13 previously produced responses between 500 - 1000 nA (Figure 1F), this suggested a change in the efficiency of receptor function, likely due to oocyte batch variability. To account for this, current responses when LEV-1 is omitted are normalized to their respective dimer control conditions (Figure 2D).

**Figure 2.**
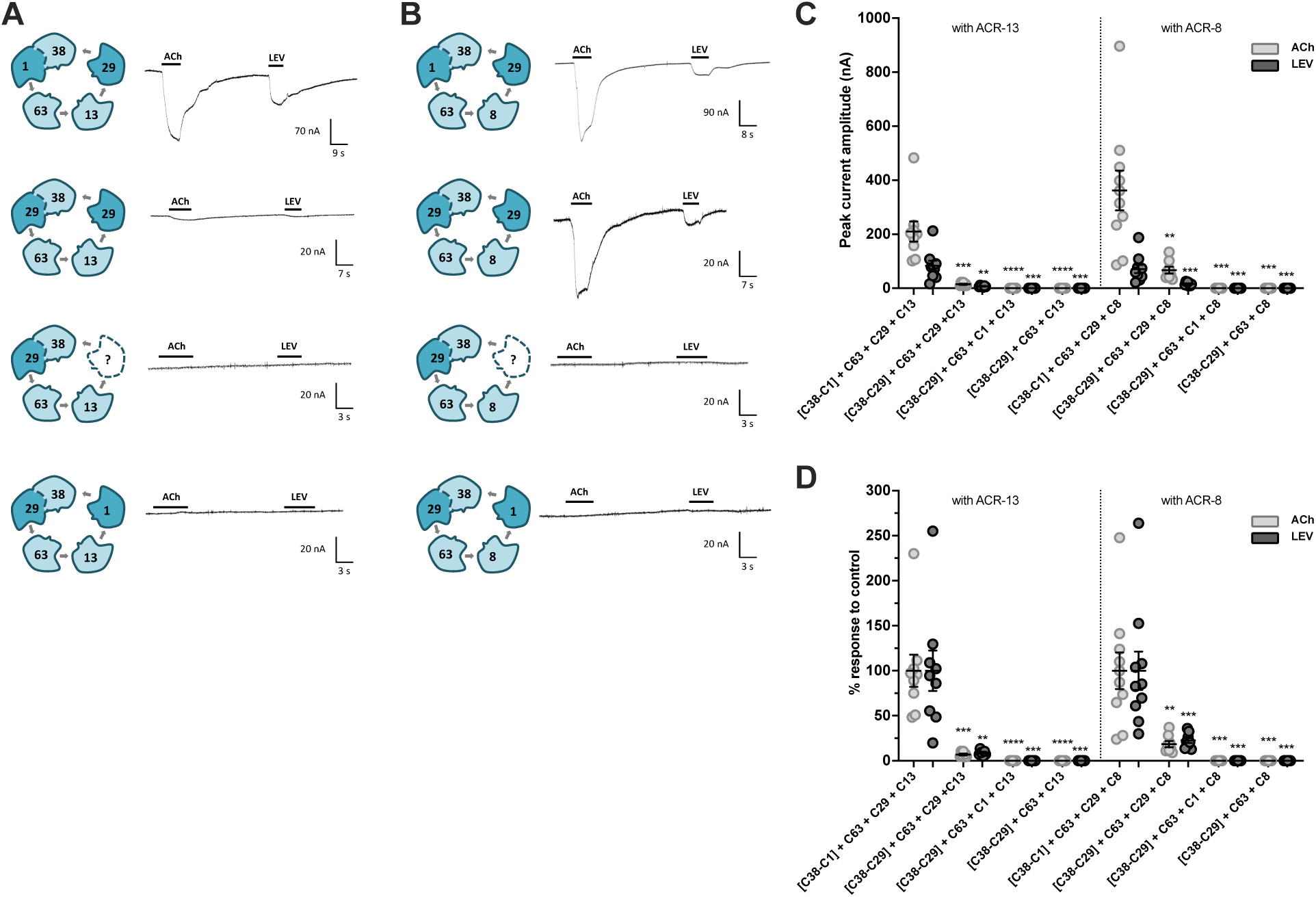
UNC-29 replaces LEV-1 in the *C. elegans* levamisole-sensitive acetylcholine receptor. A series of dimer concatemers + monomer conditions were used to identify the subunit that replaces LEV-1 when ACR-8 replaces ACR-13 in the Cel-L-AChR. (A) Sample recordings of the dimer concatemers with ACR-13 show that UNC-29 cannot replace LEV-1 with ACR-13. (B) Sample recordings of the dimer concatemers with ACR-8 show that UNC-29 replaces LEV-1 with ACR-8. (C) Summary of dimer concatemer + monomer combinations peak current amplitudes. “C” corresponds to the abbreviation of *C. elegans* followed by the subunit gene number indicated. Brackets represent subunit concatemers. Peak currents of the conditions with ACR-13 for [C38-C1] + C63 + C13 + C29 were 209.9 ± 37.42 nA, n=9 and 83.23 ± 18.68 nA, n=9; for [C38-C29] + C63 + C13 + C29 were 14.59 ± 2.06 nA, n=7 and 7.51 ± 0.9 nA, n=7; for [C38-C29] + C63 + C13 + C1 were 0 ± 0 nA, n= 10 and 0 ± 0 nA, n=10; and for [C38-C29] +C63 + C13 were 0 ± 0 nA, n= 10 and 0 ± 0 nA, n=10; to ACh and LEV, respectively. Peak currents of the conditions with ACR-8 for [C38-C1] + C63 + C8 + C2 were 361.9 ± 73.63 nA, n=10 and 71.15 ± 15.14 nA, n=10; for [C38-C29] + C63 + C8 + C29 were 66.9 ± 12.38 nA, n=8 and 16.08 ± 2.12 nA, n=8; for [C38-C29] + C63 + C8 + C1 were 0 ± 0 nA, n= 8 and 0 ± 0 nA, n=8; and for [C38-C29] + C63 + C8 were 0 ± 0 nA, n=8 and 0 ± 0 nA, n=8; to ACh and LEV, respectively. Peak current amplitudes in response to 100 μM ACh (light grey) and 100 μM LEV (dark grey). Error bars indicate SE. (D) Normalized dimer concatemer + monomer combinations. Normalized to [C38-C1] + C63 + C13 + C29, percent responses for [C38-C29] + C63 + C29 +C13 were 6.95 ± 1.0 %, n=7 and 9.03 ± 1.08 %, n=7, for [C38-C29] + C63 + C1 +C13 were 0.0 ± 0.0 %, n=10 and 0.0 ± 0.0 %, n=10; and for [C38-C29] + C63 + C13 were 0.0 ± 0.0 %, n=10 and 0.0 ± 0.0 %, n=10, for ACh and LEV respectively. Normalized to [C38-C1] + C63 + C8 + C29, percent responses for [C38-C29] + C63 + C29 +C8 were 18.49 ± 3.42 %, n=8 and 22.59 ± 2.98 %, n=9, for [C38-C29] + C63 + C8+C1 were 0.0 ± 0.0 %, n=8 and 0.0 ± 0.0 %, n=8; and for [C38-C29] + C63 + C8 were 0.0 ± 0.0 %, n=8 and 0.0 ± 0.0 %, n=8, for ACh and LEV respectively. Significance determined according to an unpaired Student’s t-test with a P-value **** p<0.0001. Error bars indicate SE. Concatenated subunits are represented in form of two subunits joined together. The number corresponds to the subunit gene name number, example UNC-38 is the subunit with the number 38. The C refers to *C. elegans*. *C. elegans* non-alpha subunits shown as dark blue and *C. elegans* alpha subunits shown as light blue. Dashed line represents the limit between the two subunits. Monomers are showed as individual subunits. See text and Supplemental Figure 1 for concatemer engineering methods. Electrophysiology traces are representative of a single oocyte recording with horizontal bars showing the time period of agonist application.

When compared to [UNC-38:LEV-1] + UNC-63 + UNC-29 + ACR-13, replacing [UNC-38:LEV-1] with [UNC-38:UNC-29] produced minimal current in response to ACh (6.95% ± 1.0 %, n = 7) and LEV (9.03 ± 1.08%, n = 7) (Figure 2A). Replacing [UNC-38:LEV-1] with [UNC-38:UNC-29] with free monomers UNC-63, UNC-29 and ACR-8 produced a more robust functional channel in response to ACh (18.49 ± 3.42 %, n = 8) and LEV (22.59 ± 2.98 nA, n = 8) (Figure 2B), confirming that UNC-29 can replace LEV-1 in the presence of ACR-8. However, these current responses represent merely ∼20% of peak currents obtained with LEV-1 indicating a preference for LEV-1 between UNC-38 and UNC-63 in the receptor. The small currents obtained with the [UNC-38:UNC-29] dimer in the presence of ACR-13 were reminiscent of some efforts to reconstitute the L-AChR with missing subunits and are likely an artefact of the *ex vivo* expression system (Fleming et al., 1997; Boulin et al., 2008). Recording traces with ACR-8 (Figure 2B, second recording) showed distinct current peaks compared to the same condition with ACR-13 (Figure 2A, second recording) which only showed small gradual changes in current upon agonist exposure. It may be that the physical link retaining UNC-29 with UNC-38 partly bypasses the normal assembly process, allowing very few receptors or partly non-functional receptors to the oocyte membrane surface with ACR-13. No receptor response was measured when the UNC-29 monomer was removed from the [UNC-38:UNC-29] + UNC-63 + ACR-8 condition (ACh 0.0 ± 0.0 %, n = 8, or LEV 0.0 ± 0.0 %, n = 8) (Figure 2B, third recording), meaning that UNC-29 is indeed present twice. This also confirms that when the UNC-29 subunit is concatenated in a dimer (ie. physically linked to UNC-38), it is unable to contribute independently to a different receptor.

Since UNC-29 is able to replace LEV-1, we wanted to test if the alternate is possible; can LEV-1 replace UNC-29 in the receptor with ACR-8? Dimer [UNC-38:UNC-29] co-injected with monomers UNC-63 + LEV-1 and ACR-8 produced no current response to ACh (0.0 ± 0.00 %, n = 8) or LEV (0.0 ± 0.00 %, n = 8) (Figure 2B, fourth recording). The same was measured when this dimer was combined with UNC-63 + LEV-1 and ACR-13 (ACh (0.0 ± 0.00 %, n = 10) and LEV (0.0 ± 0.00 %, n = 10), Figure 2A, fourth recording). To summarize, UNC-29 can physically replace LEV-1 in between UNC-38 and UNC-63 in the presence of ACR-8 and can therefore occupy both non-alpha positions in the receptor, LEV-1 is not able to replace UNC-29.

### 3.3 Functional divergence of ACR-8 and ACR-13, and among UNC-29 subunits

#### 3.3.1 Functional divergence of ACR-8 and ACR-13

A defined subunit arrangement of the Cel-L-AChR-1 and confirmation that UNC-29 replaces LEV-1 allow for accurate interpretation of receptor admixtures. Alpha subunit genes *acr-8* and *acr-13* derive from a duplication after the divergence of the Clade I nematodes. *Acr-13* was then lost in the trichostrongylid nematodes (Duguet et al., 2016; Blanchard et al., 2018). *C. elegans* ACR-8 relaxes subunit stoichiometry and allows UNC-29 to function as both non-alpha subunits, however it is not known if this relaxation is a general feature of all nematode ACR-8 subunits since their divergence from ACR-13. To test this, a series of admixtures between *C. elegans* and its relative, *H. contortus*, were made to further evaluate the functional divergence between ACR-8 and ACR-13, with a focus on Hco-ACR-8. Agreeing with previous reports ACR-13 can be replaced with ACR-8 from either *C. elegans* or *H. contortus* and produces functional receptors (Figure 3A) (Blanchard et al., 2018). Current responses for replacement with Cel-ACR-8 were ACh (387.6 ± 34.37 nA, n = 10) and LEV (80.90 ± 10.97 nA, n = 10 and for replacement with Hco-ACR-8 were ACh (955.5 ± 108.4 nA, n = 9) and LEV (1008.8 ± 113.3, n = 9) (Figure 3A and C). Removal of LEV-1 from the receptor in the presence of Hco-ACR-8 still produced ACh (383.9 ± 51.17 nA, n = 10) and LEV (420.7 ± 62.23 nA, n = 13) evoked currents, but not when UNC-29 was removed (n = 13) (Figure 3A and C). Therefore, regardless of the species of ACR-8, in the *C. elegans* receptor, UNC-29 is indispensable while LEV-1 is not.

**Figure 3.**
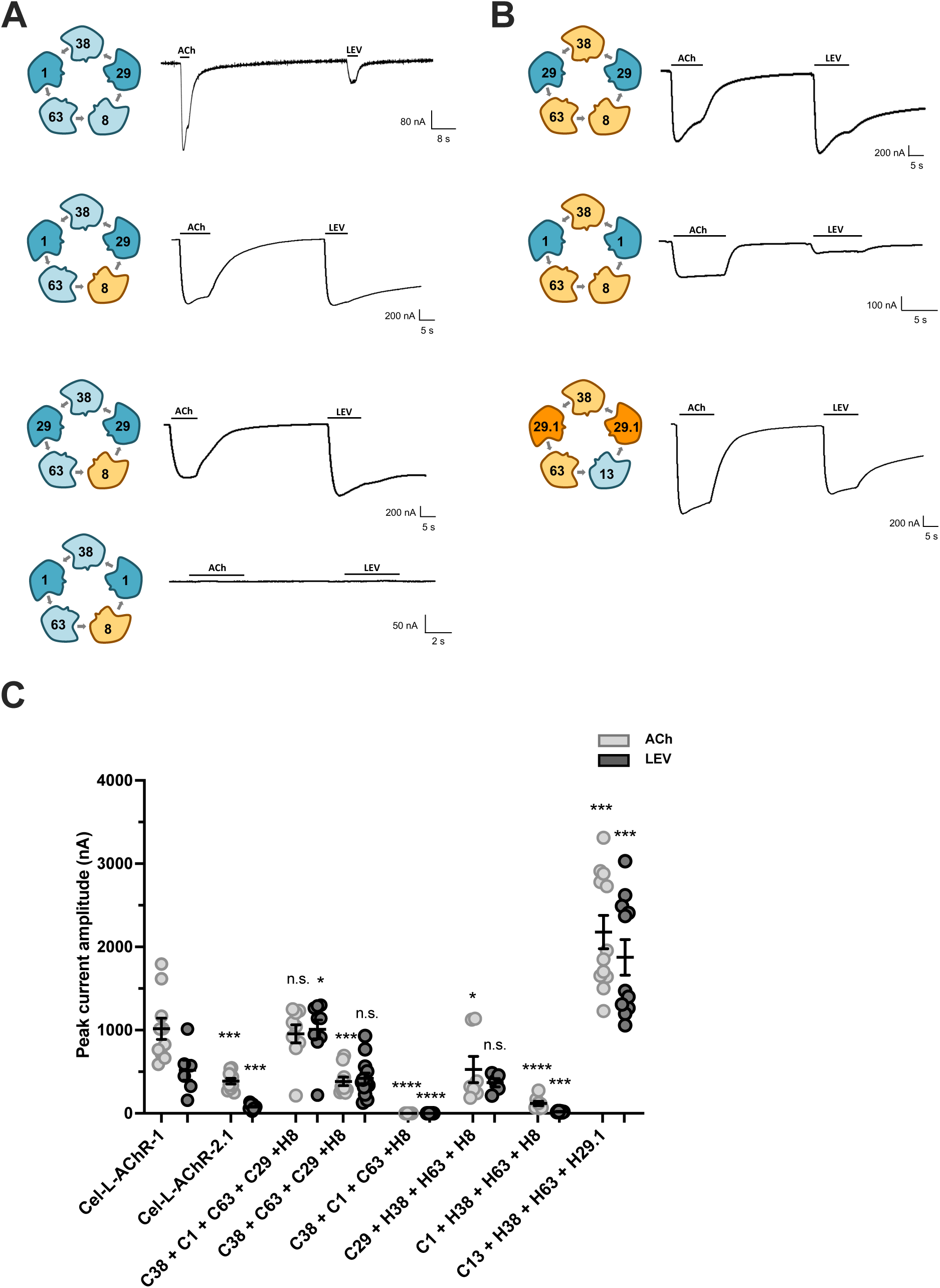
ACR-8, but not ACR-13, relaxes non-alpha subunit compatibility within the L-AChR. Admixtures between *C. elegans* and *H. contortus* subunits were used to examine the functional divergence of ACR-8 and ACR-13. (A) Sample recordings of the admixtures in a *C. elegans* receptor background. In this case, when ACR-8 replaces ACR-13, non-alpha subunit specificity becomes relaxed and produces functional receptors with only one non-alpha subunit type (UNC-29). (B) Sample recordings of the admixtures in an *H. contortus* receptor background. In this case, either UNC-29 or LEV-1 can act as the only non-alpha subunits and ACR-13 no longer requires two different non-alpha subunits. When the heteromeric receptor subunits undergo gene loss or duplication events, subunit specific is therefore relaxed allowing subunits to occupy multiple positions in the complex. (C) Summary of admixture combinations peak current amplitudes. “C” and “H” correspond to the abbreviations of *C. elegans* and *H. contortus* respectively, followed by the subunit gene number indicated. Peak currents for L-AChR-1 (with ACR13) was 1017.0 ± 128.6 nA, n=10 and 518.8 ± 118.9 nA, n=6; L-AChR-2.1 (with ACR-8) was 387.6 ± 34.4 nA, n=10, and 80.90 ± 10.97 nA, n=10; C38 + C1 + C63 + C29 + H8 was 955.5 ± 108.4 nA, n=9 and 1009.0 ± 113.3 nA, n=9; C38 + C63 + C29 + H8 was 383.9 ± 51.2 nA, n=10 and 420.7 ± 63.2 nA, n=13; C38 + C1 + C63 + H8 was 0 ± 0 nA, n=13 and 0 ± 0 nA, n=13; C29 + H38 + H63 + H8 was 527.2 ± 157.9 nA, n=7 and 368.1 ± 49.8 nA, n=5; C1 + H38 + H63 + H8 was 119.4 ± 25.6 nA, n=8 and 19.8 ± 2.5 nA, n=8; C13 + H38 + H63 + H29.1 was 2179.0 ± 200.1 nA, n=12 and 1875.0 ± 213.3 nA, n=11; for ACh and LEV, respectively. Responses were compared to the native *C. elegans* L-AChR-1 (with ACR-13). n.s.: non-significantly different according to an unpaired Student’s t-test with a P-value p<0.05. * p< 0.05; **p<0.005; ***p<0.0002, **** p<0.0001. Error bars indicate SE. Peak current amplitudes in response to 100 μM ACh (light grey) and 100 μM LEV (dark grey). Admixtures are shown as combinations of individual subunits. The number corresponds to the subunit gene name number, example UNC-38 is the subunit with the number 38. *C. elegans* non-alpha subunits shown as dark blue, *C. elegans* alpha subunits shown as light blue, *H. contortus* alpha subunits shown as light orange, and *H. contortus* non-alpha subunits shown as dark orange. Electrophysiology traces are representative of a single oocyte recording with horizontal bars showing the time period of agonist application.

Since the native *H. contortus* L-AChR does not require LEV-1 (Boulin et al., 2011) and ACR-13 is not encoded in its genome, it is easy to speculate that its L-AChR subunits have undergone relaxation allowing subunit combinations not observed with *C. elegans* subunits. To confirm this, we made admixtures in an *H. contortus* L-AChR background containing *C. elegans* subunits (Figure 3B). In the *H. contortus* receptor background, currents can be measured when Cel-UNC-29 replaces Hco-UNC-29.1 (ACh = 527.2 ± 157.9 nA, n = 7, and LEV = 369.1 ± 49.83 nA, n = 5) (Figure 3B and C). Interestingly, currents are also measured when Cel-LEV-1 is the only non-alpha subunit (ACh = 119.4 ± 25.59 nA, n = 8, and LEV = 19.82 ± 2.51 nA, n = 8) (Figure 3B and C). This indicates that LEV-1 can only act as the sole non-alpha subunit when all alpha subunits derive from *H. contortus*. This suggests that the loss of alpha ACR-13 coincides with a relaxation of subunit specificity conferred by the UNC-38 and UNC-63 subunits in the *H. contortus* receptor, allowing subunit configurations that were not previously functional to now produce functional receptors. If true, then replacing Hco-ACR-8 in the *H. contortus* receptor with Cel-ACR-13 may still produce functional channels, despite only having one non-alpha subunit type. This was indeed the case as robust currents were measured to both ACh (2179 ± 200.1 nA, n = 12) and LEV (1875 ± 213.3 nA, n = 11) when Cel-ACR-13 replaced ACR-8 in the *H. contortus* receptor (Figure 3B and C). In summary, the replacement of ACR-13 with Hco-ACR-8 removes the requirement for LEV-1 in the *C. elegans* receptor. However, in receptors where multiple instances of gene loss and duplication occur, then subunit specificity becomes relaxed, allowing subunits to occupy multiple positions/roles to maintain a pentamer.

#### 3.3.2 Functional divergence of *H. contortus* UNC-29.2

The trichostrongylid nematode *unc-29* subunits have duplicated multiple times, with *H. contortus* having four full-length copies (Duguet et al., 2016; Doyle et al., 2020). Additionally, although *H. contortus* has a *lev-1* gene predicted in its genome, there is no signal peptide or any evidence that it contributes to the L-AChR *in vivo* (Neveu et al., 2010; Boulin et al., 2011). Previous work on the characterization of the four UNC-29 copies found that while all are functional, UNC-29.2 is the only one that absolutely requires LEV-1, the others do not (Duguet et al., 2016). The results of our ACR-8 and ACR-13 admixtures suggest that gene duplication and loss coincide with changes to subunit composition and compatibility. A series of admixtures were therefore made to further investigate the functional divergence of UNC-29.2. As shown previously all Hco-UNC-29 paralogs are functional in the *C. elegans* receptor when including LEV-1 (UNC-38 + LEV-1 + UNC-63 + ACR-13 + Hco-UNC-29.1 (ACh = 471.1 ± 51.48 nA, n = 9, and LEV = 279.9 ± 34.02 nA, n = 11), UNC-38 + LEV-1 + UNC-63 + ACR-13 + Hco-UNC-29.2 (ACh = 28.61 ± 5.17 nA, n = 10, and LEV = 16.87 ± 3.83 nA, n = 10), UNC-38 + LEV-1 + UNC-63 + ACR-13 + Hco-UNC- 29.3 (ACh = 109.6 ± 20.88 nA, n = 9, and LEV = 38.93 ± 7.03 nA, n = 12), and UNC-38 + LEV-1 + UNC-63 + ACR-13 + Hco-UNC-29.4 (ACh = 49.20 ± 14.50 nA, n = 5, and LEV = 14.63 ± 2.64 nA, n = 6)) (Figure 4A) (Duguet et al., 2016). Removal of LEV-1 abolished responses with Hco- UNC-29.2 (UNC-38 + UNC-63 + ACR-13 + Hco-UNC-29.2 (n=8) but not with the other paralogs ((UNC-38 + UNC-63 + ACR-13 + Hco-UNC-29.1 (ACh 1766.05 ± nA =, n=10, and LEV = 1637.3 ± nA, n=10), UNC-38 + UNC-63 + ACR-13 + Hco-UNC-29.3 (ACh = 337.4 ± 72.86 nA, n = 9, and LEV = 73.24 ± 15.51 nA, n = 14) and UNC-38 + UNC-63 + ACR-13 + Hco-UNC-29.4 (ACh = 346.9 ± 84.10 nA, n = 6, and LEV = 101. 3 ± 31.80 nA, n = 11)) (Figure 4A). A summary of the responses is shown in Figure 4B. UNC-29.2 is therefore inherently functional but requires the addition of the non-alpha LEV-1 subunit.

**Figure 4.**
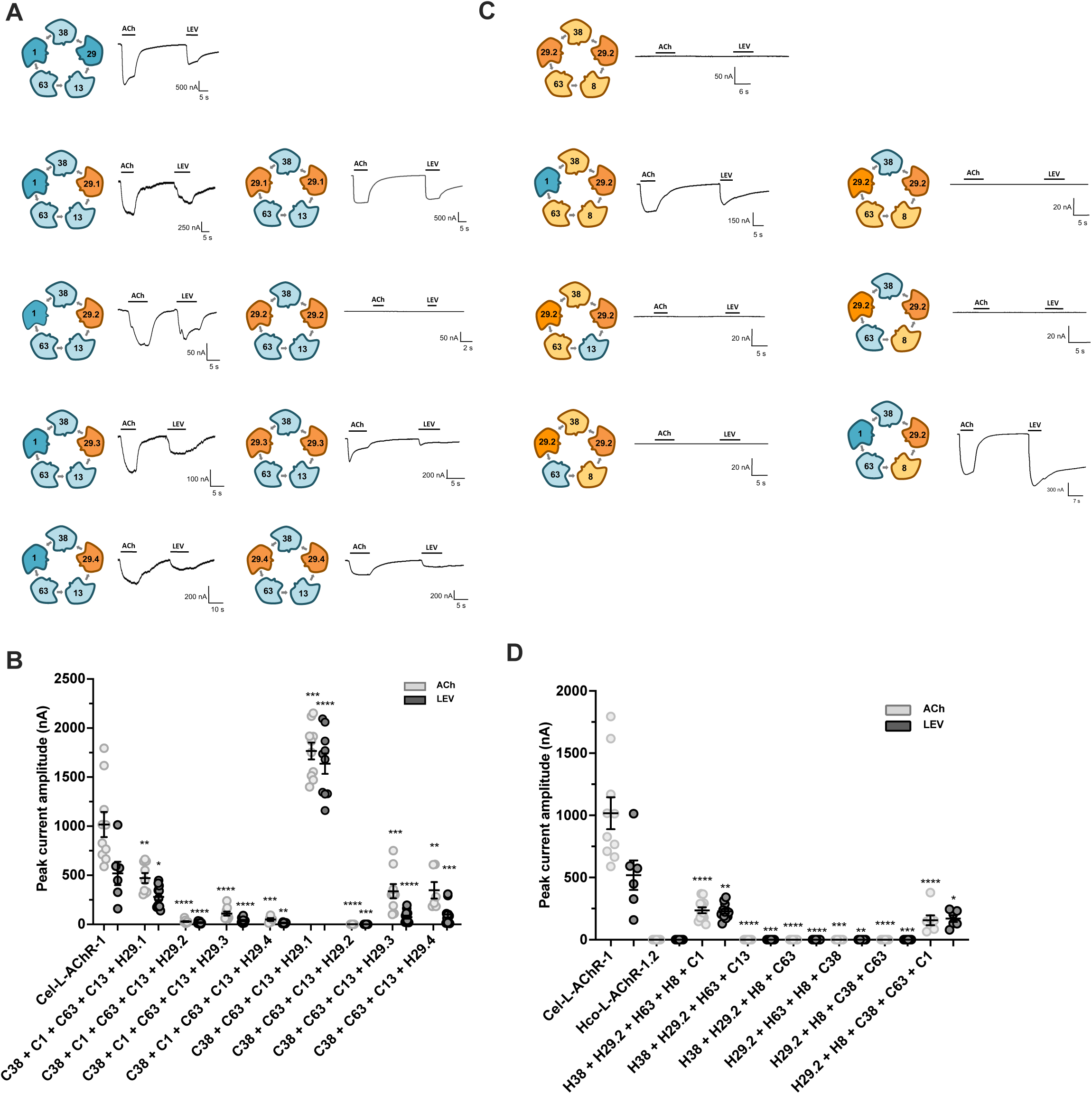
*H. contortus* UNC-29.2 cannot function as the only non-alpha subunit in the L-AChR. Admixtures between *C. elegans* and *H. contortus* subunits were used to examine the functional divergence of Hco- unc-29.1and Hco-unc-29.2. (A) Sample recordings of the admixtures show that all Hco-unc-29 paralogs are functional and can replace Cel-UNC-29 the *C. elegans* L-AChR. However, upon removal of LEV-1, Hco-UNC-29.2 no longer produces a functional channel whereas the other Hco-unc-29 paralogs can. Hco-unc-29.2 cannot function as the only non-alpha subunit. (B) Summary of admixture combinations peak current amplitudes. “C” and “H” correspond to the abbreviations of *C. elegans* and *H. contortus* respectively, followed by the subunit gene number indicated. Peak currents for L-AChR-1 was 1017.0 ± 128.6 nA, n=10 and 518.8 ± 118.9 nA, n=6; C38 + C1 + C63 + C13 + H29.1 was 471.1 ± 51.5 nA, n=9 and 279.9 ± 34.0 nA, n=11; C38 + C1 + C63 + C13 + H29.2 was 28.6 ± 5.2 nA, n=10 and 16.9 ± 3.8 nA, n=9; C38 + C1 + C63 + C13 + H29.3 was 109.6 ± 20.9 nA, n=9 and 38.8 ± 7.0 nA, n=12; C38 + C1 + C63 + C13 + H29.4 was 49.2 ± 14.5 nA, n=5 and 14.6 ± 2.6 nA, n=6; C38 + C63 + C13 + H29.1 was 1766.1 ± 85.6 nA, n=10 and 1637.3 ± 103.9 nA, n=10;C38 + C63 + C13 + H29.2 was 0 ± 0 nA, n=8 and 0 ± 0 nA, n=8; C38 + C63 + C13 + H29.3 was 337.4 ± 72.9 nA, n=9 and 73.2 ± 15.6 nA, n=14; C38 + C63 + C13 + H29.4 was 346.9 ± 84.1 nA, n=6 and 101.3 ± 31.8 nA, n=11; for ACh and LEV, respectively. Current responses of *H. contortus* unc-29 subunits in the *C. elegans* L-AChR with LEV-1 were taken from Duguet et al., 2016. Responses were compared to the native *C. elegans* L-AChR-1 (with ACR-13). n.s.: non-significantly different according to an unpaired Student’s t-test with a P-value > 0.05. * p< 0.05; **p<0.01; ***p<0.005, **** p<0.0001. Error bars indicate SE. Peak current amplitudes in response to 100 μM ACh (light grey) and 100 μM LEV (dark grey). (C) Sample recordings of the admixtures show that under all admixture combinations examined, Hco-unc-29.2 requires another non-alpha subunit in the receptor. (D) Summary of admixture combinations peak current amplitudes. “C” and “H” correspond to the abbreviations of *C. elegans* and *H. contortus* respectively, followed by the subunit gene number indicated. Peak currents for L-AChR-1 was 1017.0 ± 128.6 nA, n=10 and 518.8 ± 118.9 nA, n=6; Hco-L-AChR-1.2 (with UNC-29.2) was 0 ± 0 nA, n=, and 0 ± 0 nA, n=; H38 + H29.2 + H63 + H8 + C1 was 236.0 ± 23.9 nA, n=12, and 227.9 ± 17.8 nA, n=12; H38 + H29.2 + H63 + C13 was 0 ± 0 nA, n=8, and 0 ± 0 nA, n=;8 H38 + H29.2 + H8 + C63 was 0 ± 0 nA, n=10, and 0 ± 0 nA, n=10; H29.2 + H63 + H8 + C38 was 0 ± 0 nA, n=5, and 0 ± 0 nA, n=5; H29.2 + H8 + C38 + C63 was 0 ± 0 nA, n=8, and 0 ± 0 nA, n=8; H29.2 + H8 + C38 + C63 + C1 was 156.1 ± 39.8 nA, n=7, and 169.9 ± 27.1 nA, n=6; for ACh and LEV, respectively. Responses were compared to the native *C. elegans* L-AChR-1 (with ACR-13). n.s.: non-significantly different according to an unpaired Student’s t-test with a P-value p<0.05. * p< 0.05; **p<0.005; ***p<0.0002, **** p<0.0001. Error bars indicate SE. Peak current amplitudes in response to 100 μM ACh (light grey) and 100 μM LEV (dark grey). Admixtures are shown as combinations of individual subunits. The number corresponds to the subunit gene name number, example UNC-38 is the subunit with the number 38. *C. elegans* non-alpha subunits shown as dark blue, *C. elegans* alpha subunits shown as light blue, *H. contortus* alpha subunits shown as light orange, and *H. contortus* non-alpha subunits shown as dark orange. Electrophysiology traces are representative of a single oocyte recording with horizontal bars showing the time period of agonist application.

We next wanted to test the function of UNC-29.2 in a more native environment, namely with other *H. contortus* subunits. Starting with the *H. contortus* receptor with UNC-29.2 which produces no response to ACh or LEV (n = 9) (Duguet et al., 2016), we replaced each subunit individually with its *C. elegans* ortholog and measured responses as before (Figure 4C). Only when LEV-1 was added to the admixture was a receptor current obtained in response to ACh (236.0 ± 23.89 nA, n = 12) and LEV (227.9 ± 17.76 nA, n = 12) (Figure 4C). Replacement of Hco-ACR-8 with Cel-ACR- 13 (n = 8), Hco-UNC-63 with Cel-UNC-63 (n=10), and Hco-UNC-38 with Cel-UNC-38 (n = 5) all failed to produce functional receptors (Figure 4C).

In Figure 3B, we found that replacing Cel-UNC-38 and Cel-UNC-63 with their *H. contortus* orthologs allowed LEV-1 to act as the only non-alpha subunit. Therefore, we exchanged Hco- UNC-38 together with Hco-UNC-63 for their *C. elegans* orthologs to observe if UNC-29.2 could produce a measurable channel under these circumstances. No response was measured for either agonist (Hco-UNC-29.2 + Hco-ACR-8 + Cel-UNC-38 + Cel-UNC-63, n = 8) (Figure 4B). However, adding LEV-1 to this admixture did produce a functional channel (Hco-UNC-29.2 + Hco-ACR-8 + Cel-UNC-38 + Cel-UNC-63 + Cel-LEV-1 for ACh was 156.1 ± 39.80 nA, n = 7, and for LEV was 169.9 ± 27.07 nA, n = 6) (Figure 4B). This demonstrates that regardless of the L-AChR background (*C. elegans* or *H. contortus)* or ACR-8 or ACR-13, Hco-UNC-29.2 requires another non-alpha subunit to contribute to a function channel. A summary of the responses is shown in Figure 4D.

### 3.4 The intracellular loop determines non-alpha subunit placement within the receptor

The functional difference between Hco-UNC-29.1 and Hco-UNC-29.2 must have occurred since their duplication and would be encoded in their sequences. Their amino acid sequences differ by approximately 25%, with the majority of differences found in the intracellular loop (ICL), a region involved in receptor expression (Rudell et al., 2014, 2020). To identify the general region that mediates the difference between the two subunits, chimeras were made exchanging the ICL between *unc-29.1* and *unc-29.2,* as shown in Figure 5A. Chim-1 has the extracellular and transmembrane domain sequences of *unc-29.1* with the ICL sequence of *unc-29.2*, whereas Chim-2 has the extracellular and transmembrane domain sequences of *unc-29.2* sequence with the ICL sequence of *unc-29.1*. These chimeras were expressed with combinations of other subunits to evaluate their requirement for the LEV-1 subunit.

**Figure 5.**
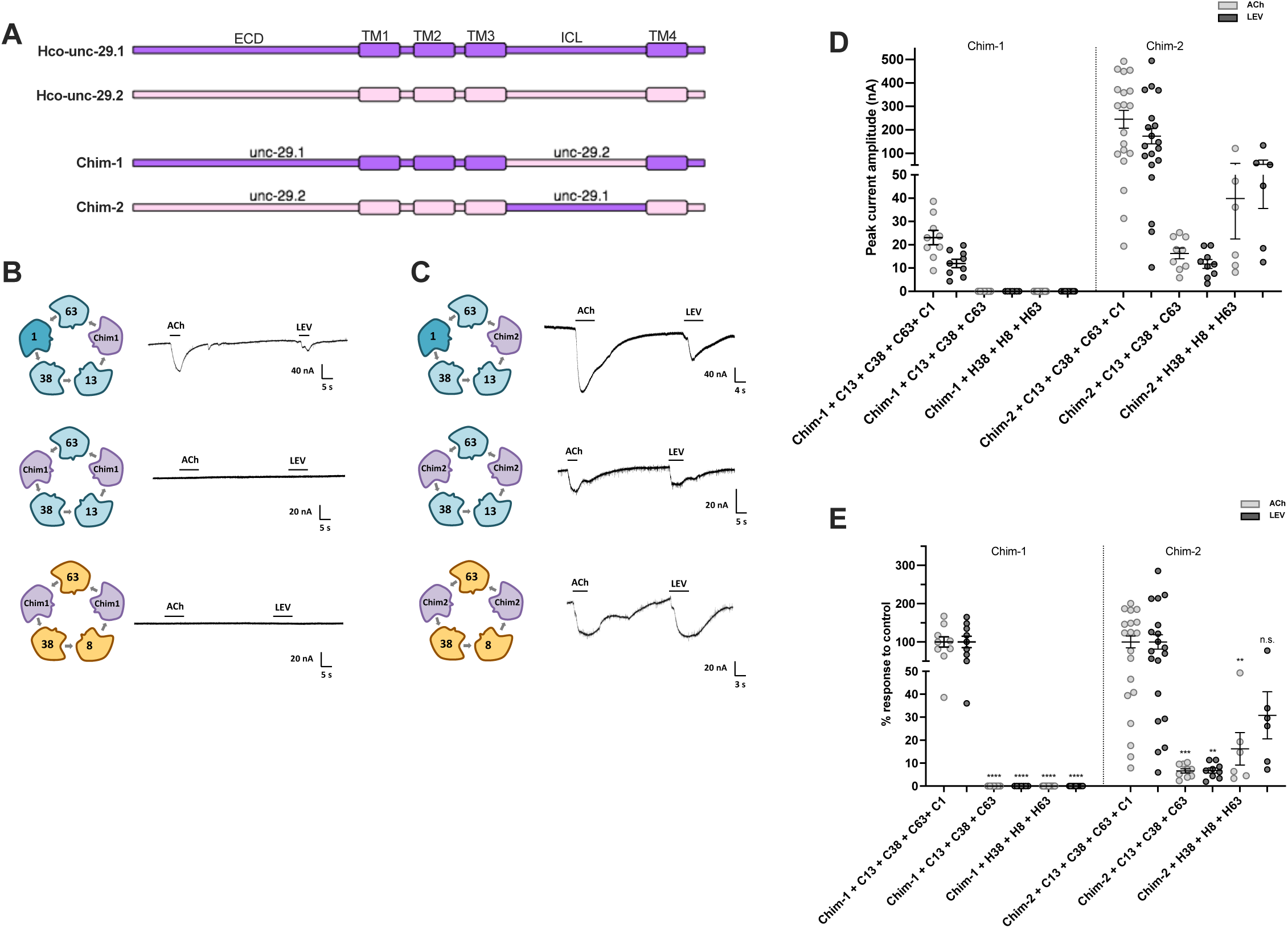
The intracellular loop determines unc-29 positional plasticity. Chimeras were made between *H. contortus unc-29.1* and *unc-29.2* by exchanging the intracellular loop to identify the region mediating the ability of unc-29.1 to function as the only non-alpha subunit in the receptor. (A) Chim-1 has the *unc-29.1* sequence with the intracellular loop of *unc-29.2*, and Chim-2 has the *unc-29.2* sequence with the intracellular loop of *unc-29.1*. (B) Sample recordings of the Chim-1 conditions show that despite the chimera being functional (second recording), it cannot produce current responses in oocytes without LEV-1 in either the Cel-LAChR background (third recording) or the Hco-L-AChR background (bottom recording). (C) Sample recordings of the Chim-2 conditions show that the chimera is functional (top recording) and that it remains functional without LEV-1 in both the Cel-LAChR background (middle recording) and the Hco- L-AChR background (bottom recording). Chim-2 produced measurable responses without Lev-1, therefore the ICL mediates the ability of UNC-29.1 to function as the only non-alpha subunit and occupy two positions in the receptor. (D) Summary of unc-29 chimera peak current amplitudes. “C” and “H” correspond to the abbreviations of *C. elegans* and *H. contortus* respectively, followed by the subunit gene number indicated. Chim-1 produced small but measurable responses to both ACh (23.1 ± 3.1 nA, n=9) and LEV (11.9 ± 1.8 nA, n=9) in the Cel-L-AChR with LEV-1. No current was measured in the absence of LEV-1 in both the *C. elegans* (n=7) and *H. contortus* (n=11) L-AChR subunits. Chim-2 produced measurable responses to both ACh (245.5 ± 37.8 nA, n=19) and LEV (173.2 ± 32.6 nA, n=19) in the Cel-L-AChR with LEV-1. Small currents were measured in the absence of LEV-1 in both the *C. elegans (*16.27 ± 2.3 nA and 11.74 ± 1.9 nA, n=9) and *H. contortus* (39.87 ± 1.9 nA and 53.3 ± 17.8 nA, n=6) L-AChR subunits, for ACh and LEV respectively. (E) Normalized responses for chimeras from Panel C. Chim-1 no longer produced responses to ACh (0.00 ± 0.00 %, n=7) or LEV (0.00 ± 0.00 %, n=7) in the *C. elegans* receptor background or in the *H. contortus* receptor background (0.00 ± 0.00 % for both ACh and LEV, n=11). Chim-2 produced small yet measurable responses to ACh (6.63 ± 0.9 %, n=9) and LEV (6.78 ± 1.1 %, n=9) in the *C. elegans* subunit background without LEV-1. Larger sized responses were measured in the *H. contortus* receptor background for ACh (16.24 ± 7.08 %, n=6) and LEV (30.79 ± 10.25 %, n=6). n.s.: non-significantly different according to an unpaired Student’s t-test with a P-value p<0.05. **p<0.005; ***p<0.0002, **** p<0.0001. Error bars indicate SE. Chimera admixtures are shown as combinations of individual subunits. The number corresponds to the subunit gene name number, example UNC-38 is the subunit with the number 38. The C corresponds to *C. elegans*. *C. elegans* non-alpha subunits shown as dark blue, *C. elegans* alpha subunits shown as light blue, *H. contortus* alpha subunits shown as light orange, and Chimeras shown as purple. Electrophysiology traces are representative of a single oocyte recording with horizontal bars showing the time period of agonist application.

Subunit combinations in the Cel-LAChR-1 where the chimeras simply replaced UNC-29 served as a control to ensure that exchanging the ICL did not render the chimeras non-functional (Figure 5 B & C). Chim-1 example recordings shown in Figure 5B, Chim-2 example recordings shown in Figure 5C, with summary responses in Figure 5D. Chim-1 responses to ACh were (23.07 ± 3.08 nA, n = 9) and to LEV were (11.94 ± 1.78 nA, n = 9) and Chim-2 responses to ACh (245.5 ± 37.84 nA, n = 19) and LEV (173.2 ± 32.58 nA, n = 19). The smaller response measured with Chim-1 and large variability observed with Chim-2 may be due to inefficient receptor expression introduced by exchanging their ICLs. However, both chimeras produced consistent and measurable currents in response to both ligands and their values were within range of those obtained with the C*. elegans* L-AChR containing Hco-UNC-29.1 or Hco-UNC-29.2 (Figure 4A, left). Therefore the chimeras functioned comparably to their “parent” subunits. To account for the differences in response sizes between the two chimeras, chimera combinations where LEV-1 was not included are shown relative to the responses obtained when LEV-1 is present (Figure 5E). Chim-1 no longer produced responses to ACh (0.00 ± 0.00 %, n = 7) or LEV (0.00 ± 0.00 %, n = 7) in the *C. elegans* receptor background when LEV-1 was omitted. The same was observed in the *H. contortus* receptor background (0.00 ± 0.00 % for both ACh and LEV, n = 11). Chim-2 on the other hand produced small yet measurable relative responses to ACh (6.63 ± 0.9 %, n = 9) and LEV (6.78 ± 1.1 %, n = 9) in the *C. elegans* subunit background without LEV-1. Larger sized relative responses were measured in the *H. contortus* receptor background for ACh (16.24 ± 7.08 %, n = 6) and LEV (30.79 ± 10.25 %, n = 6), likely because the subunits are from *H. contortus*. Chim-1 behaved as UNC-29.2, requiring LEV-1 in either the *C. elegans or H. contortus L-AChRs* and Chim-2 behaved as UNC-29.1, not requiring LEV-1 in either L-AChR. These results indicate that the ICL is responsible for most of the phenotype difference and is therefore a region that contributes to non-alpha subunit compatibility and positioning. However, it is important to note that the smaller responses obtained with chim-2 without LEV-1 indicate problems with receptor function. It could be that another region in combination with the ICL is involved. Despite this, the fact that the ICL exchange completely ablated UNC-29.1 response and allowed UNC-29.2 to produce a functional channel confirms that the ICL is a major region.

## Discussion

The numerous pLGIC subunit gene duplication and loss events in the nematodes gave rise to an expanded receptor family with complex receptor possibilities (Dent, 2006; Jones and Sattelle, 2008; Beech and Neveu, 2015). Since the functional properties of a channel depend on its composition and organization, knowing the mechanisms that underly this would provide valuable insight into the constraints on functional evolutionary change of this important drug target. Despite decades of research on the L-AChR, its stoichiometry and the mechanisms determining subunit compatibility remain unknown. The goal of this study was to therefore to identify the subunit stoichiometry of the *C. elegans* L-AChR and further elucidate the functional divergence of alpha subunits ACR-8 and ACR-13 and non-alpha subunits UNC-29.1 and UNC-29.2.

A number of methods exist to identify subunit positioning within pLGICs (Nashmi et al., 2003; Minier and Sigel, 2004; Ericksen and Boileau, 2007; Millar, 2009; Hanson and Czajkowski, 2011). The approach used here was to link subunits together into one continuous construct. The C-terminal tail of the first subunit was linked to the N-terminal domain of the next subunit. Only the first subunits’ signal peptide and the last subunits’ stop codon were retained, making a single open reading frame and restricting the subunits to one specific configuration. This technique is commonly used when determining subunit stoichiometry (Zhou et al., 2003; Groot-Kormelink et al., 2006; Carbone et al., 2009). However, linker length must be optimized because short linkers can introduce steric hindrance and long linkers can allow other subunits to be inserted between the two linked subunits (Liao et al., 2020, 2021). Computer modelling of the dimer concatemers used here confirmed that the distance between them did not impose steric hindrance nor allow insertion of other subunits from occurring (Supplemental Figure 2). Only the UNC-63 C-terminal tail required the addition of an artificial linker to ensure sufficient length to connect it to the next subunit. This linker had no effect on the function of the UNC-63 subunit (Figure 1F). Our concatemers produced functional receptors, so the technique did not block function by steric hindrance. However, reduced responses were observed as the concatemer length increased. Regardless, this technique offered direct manipulation of the subunits for testing specific arrangements.

An initial list of dimeric concatemers was made to begin with combinations where the first subunit was an alpha-type. This reduced the number of combinations to test and guaranteed that at least three would have subunits in the right order. Unexpectedly, only the [UNC-38:LEV-1] dimer produced measurable current response (Figure1A). Therefore, two dimers had subunits in the correct order but failed to produce a functional channel. Either this was due to technical failure or the dimer combinations interfered with receptor assembly. The acetylcholine receptor assembly process is tightly regulated and can require the addition of subunits in a specific order (Green and Claudio, 1993; Green, 1999; Wanamaker et al., 2003).

A series of trimeric constructs were then made based on the success of the of the [UNC-38:LEV-1] dimer. Again, only one resulted in a functional channel; [UNC-38:LEV-1:UNC-63] (Figure 1B). This meant that one of the other trimers that had UNC-29 or ACR-13 before UNC-38 was in the correct order but did not produce a functional channel. This suggested that our concatemer strategy could provide information on the L-AChR assembly process in addition to the subunit stoichiometry. The sequential model of receptor assembly states that an initial trimer of three subunits is quickly formed followed by the sequential, slower addition of the two remaining subunits (Green and Claudio, 1993; Green and Wanamaker, 1997, 1998; Wanamaker et al., 2003). This model explains why no functional receptors were produced with two dimers and one trimer despite having the correct order ([UNC-63:ACR-13], [ACR-13:UNC-29] and [UNC-29:UNC- 38:LEV-1]). If true, then the [LEV-1:UNC-63] dimer that was not initially tested would produce a functional receptor. Indeed it did, but with currents lower than the [UNC-38:LEV-1] dimer, possibly due to it being the last two subunits in the trimer (Figure 1C).

Two tetramer and two pentamer concatemers were made based on the success of the [UNC- 38:LEV-1:UNC-63] trimer. The [UNC-38:LEV-1:UNC-63:ACR-13] tetramer with UNC-29 as a monomer produced a response with severely dampened currents. The other tetramer, [UNC- 38:LEV-1:UNC-63:UNC-29] with ACR-13 as a monomer, did not produce measurable responses (Figure 1D). This supports the quaternary structure of the *C. elegans* L-AChR having an UNC- 38:LEV-1:UNC-63:ACR-13:UNC-29 counterclockwise arrangement when viewed from outside the cell. An initial trimer of UNC-38:LEV-1:UNC-63 is formed followed by the sequential addition of the remaining subunits ACR-13 then UNC-29 (Figure 6C). The proposed subunit arrangement agrees with *in silico* models that predict optimal ligand orientation and binding energies in the UNC-38/LEV-1 and ACR-13/UNC-29 and interfaces (Hernando et al., 2012). Unfortunately, neither pentameric construct yielded any response, likely due to difficulties with expressing such long concatemers (Figure 1E). The large currents measured with the trimer (∼461 nA for ACh) were in contrast to the small currents observed with the tetramer (∼37 nA for ACh) and indicated that each additional subunit in the concatemer chain interfered with normal assembly, leading to dampened current response. The tetrameric and pentameric concatemers may introduce problems for the normal assembly and processing in the oocyte, as observed in vertebrate receptors (Zhou et al., 2003; Groot-Kormelink et al., 2006; Carbone et al., 2009; Shu et al., 2012). Further work is required to confirm the order of these last two subunits.

**Figure 6.**
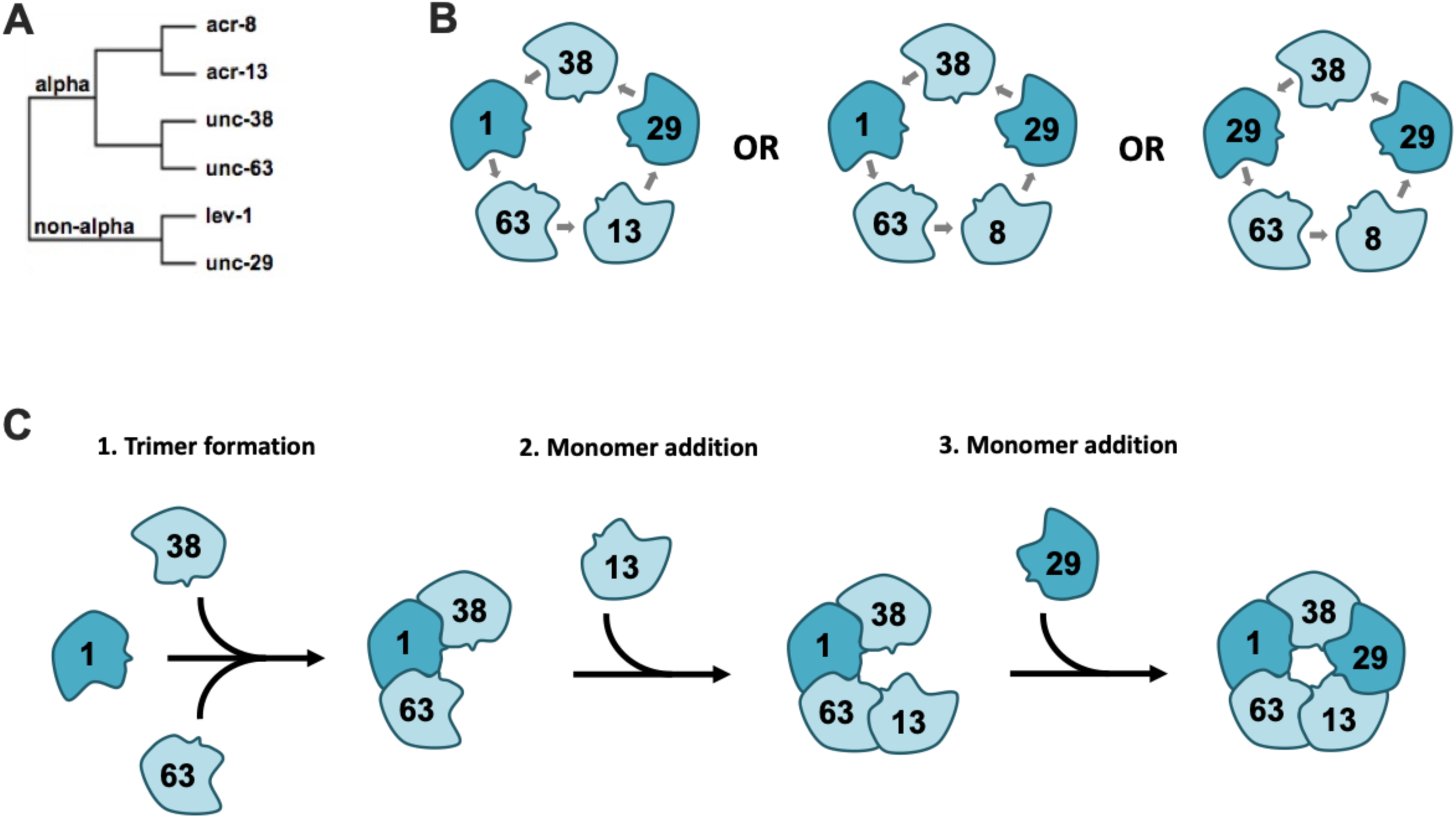
Proposed Cel-L-AChR stoichiometry and assembly. (A) Phylogeny of L-AChR subunits separates the non-alpha and alpha subunit types first, followed by further duplication and divergence into different subunits within these classes. (B) Proposed arrangement of the L-AChR separates non-alpha subunits and is, in a counterclockwise order; UNC-38-LEV-1-UNC-63-ACR-13-UNC-29. Replacement of ACR-13 with ACR-8 relaxes the requirement for five distinct subunits, LEV-1 is now dispensable and can be replaced with UNC-29. Cel- L-AChR nomenclature taken from Blanchard et al., 2018. (C) Proposed assembly pathway based on the concatemers used here supports the sequential model of receptor formation (Green and Claudio, 1993). An initial trimer is formed of UNC-38-LEV-1-UNC-63, followed by individual addition of ACR-13 and UNC-29. Non-alpha subunits in dark blue, alpha subunits in light blue.

We next evaluated the functional divergence of different subunit types using cross-species admixtures. We used admixtures from Clade V nematodes *H. contortus* and *C. elegans* to evaluate the divergence of ACR-8 & ACR-13 and UNC-29.1 & UNC-29.2. Both species of L-AChR require the same accessory proteins (RIC-3, UNC-50 and UNC-74) (Boulin et al., 2008, 2011) and are therefore under similar regulatory mechanisms.

Alpha subunit duplication gave rise to *acr-8* and *acr-13* which have since functionally diverged (Figure 6A). *C. elegans* has retained both while *H. contortus* had only retained *acr-8* (Boulin et al., 2011; Blanchard et al., 2018). Both contribute to a robust L-AChR, but ACR-8 does not require LEV-1 while ACR-13 does (Boulin et al., 2008, 2011; Blanchard et al., 2018). In this case, the specificity that prevents UNC-29 from replacing LEV-1 is determined by an interaction with ACR-13. No functional receptor was produced when LEV-1 was the only non-alpha subunit, which implies that the mechanism that confers specificity on the LEV-1 subunit remains intact even in the presence of ACR-8, from *C. elegans* or *H. contortus*.

Interestingly, the admixture containing *H. contortus* UNC-38, UNC-63 and ACR-8 with Cel-LEV-1 as the only non-alpha subunit produced a functional channel. Our admixture results suggest that heteromeric subunits become specialized for their specific roles. The *C. elegans* L-AChR with ACR-13 represents an extreme example where five different subunits are needed with each confined to a single position and function (Boulin et al., 2008). This is in contrast to homomeric subunits that must occupy all positions and roles within the receptor. Subunit duplication and loss events disrupt the evolved subunit specificity causing subunits to relax specificity requirements in order to still produce a functional channel. This may explain why receptors can be produced from different combinations of subunits from *A. suum, H. contortus,* and *O. dentatum* (Williamson et al., 2009; Boulin et al., 2011; Buxton et al., 2014) and why LEV-1 can function as the only non-alpha subunit with the *H. contortus* alpha subunits but not the *C. elegans* alpha subunits.

Our admixture conditions where LEV-1 is omitted indicate at least one other subunit is present twice. The fact that UNC-29 and LEV-1 are more closely related phylogenetically (Figure 6A) and the importance of having two non-alpha subunits in a heteromeric receptor suggests that UNC-29 likely replaces LEV-1. We carried out a series of dimer concatemer and monomer combinations to test this (Figure 2). Only very small currents were measured with the [UNC-38:UNC-29] dimer in the presence of ACR-13, likely an artefact of the expression system (Fleming et al., 1997; Boulin et al., 2008). Larger and more consistent current responses were measured when the same dimer was in combination with ACR-8. Based on our proposed model of subunit arrangement and assembly, the specificity for LEV-1 is conferred only after the initial trimer of [UNC-38:LEV-1:UNC-63] is formed. The admixture of *C. elegans* UNC-38, UNC-63 and UNC-29 with ACR-8 produced a functional channel, therefore UNC-38, UNC-29 and UNC-63 can form the initial trimer complex but it only becomes functional if ACR-8 joins afterwards, not ACR-13. The fact that the [UNC-38:UNC-29] dimer produced smaller maximal currents than [UNC-38:LEV-1] when ACR-8 was present suggests that the LEV-1 subunit in the initial trimer is preferred. Regardless, the ACR-13 subunit confers the specificity for LEV-1 after the initial trimer has formed and without directly contacting it, since they are separated by UNC-63 in the pentamer (Figure 6B). The mechanism described is likely different from the ability of Hco-UNC-29.1 to remove the requirement for LEV-1, since in each case it is different subunit types (alpha vs non-alpha) mediating the requirement of a non-alpha subunit. It is also important to note that we only tested if UNC-29 can replace LEV-1, additional dimer and monomer conditions would directly test if the alpha subunits are also able to replace LEV-1, although this is unlikely.

Our Hco-UNC-29 chimeras identified the ICL, the most variable sequence region in pLGICs, as being responsible for the ability to remove the requirement for LEV-1. However, the small percent responses obtained with Chim-2 with the *H. contortus* (∼16 %) or *C. elegans* (∼6 %) receptor backgrounds indicate inefficient receptor function. Another region, in combination with the ICL, may also be involved. However, the fact that the ICL exchange completely ablated UNC-29.1 response and allowed UNC-29.2 to produce a functional channel indicates that the ICL is a determining region. It is not currently known if UNC-29.1 is the subunit that replaces LEV-1, although this seems likely based on our concatemer results. The structure and function of the ICL remain uncharacterized. The only defined structures are two alpha helices at each end of it, with one acting as a funnel for ions crossing the channel pore (Unwin, 2005). The ICL has been found to mediate diverse functions ranging from receptor trafficking (Ren et al., 2005) to receptor kinetics (Peters et al., 2005; Hales et al., 2006; McKinnon et al., 2012). Our chimeras indicate that the ICL plays an important role in subunit stoichiometry and may make inter-subunit interactions that determine their compatibility. It may be an important region for regulating receptor diversity since the minimum sequence length required for the ICL in eukaryotes is 70 amino acids (the ICLs studied here are 150) and large, diverse ICLs emerged in animal pLGICs (Tasneem et al., 2005; Baptista-Hon et al., 2013).

Taken together, combining cross-species admixtures with subunit concatemers is an informative and robust strategy to decipher subunit stoichiometry and interactions. This method is particularly relevant to determine subunit positioning and diversity within the parasitic nematode L-AChRs. Using our proposed *C. elegans* L-AChR arrangement, and according to the different functional subunit admixtures, a similar approach of concatenation could also be applied to helminth receptors. Additionally, this approach could determine if the ACR-8 from *H. contortus* simply replaces *C. elegans* ACR-13 or if it induces a rearrangement of the other subunits and to determine the stoichiometry of the multiple Hco-UNC-29 copies. In a more general context, our results also indicate that loss or duplication of heteromeric receptor subunit genes disrupt the previously evolved subunit arrangement and inter-subunit interactions. This causes relaxation in the maintenance of a specific stoichiometry and in turn allows subunits to occupy multiple positions and functions within the receptor. Finally, the identification of the quaternary structure of the *C. elegans* L-AChR as UNC-38:LEV-1:UNC-63:ACR-13:UNC-29 represents a final step in the search for the structure of this receptor since the initial screen for LEV resistant mutants of *C. elegans* in the early 1980’s (Lewis et al., 1980b, 1980a).

## Author contributions

JDN- Conceptualization, Data curation, Formal analysis, Investigation, Methodology, Validation, Visualization, Writing – original draft, review & editing.

TBD- Conceptualization, Data curation, Formal analysis, Investigation, Methodology, Validation, Visualization, Writing – original draft, review & editing.

CJHH – Data curation and Investigation

RNB – Conceptualization, Funding, Methodology, Visualization, Writing – review.

## Acknowledgments

We thank C. Neveu for generously providing the following subunit genes in the pTB expression plasmid from C. elegans (lev-1, acr-13, unc-63a, unc-38 and unc-29) and H. contortus (unc-29.1, unc-29.2, unc-29.3, unc-29.4, unc-38, unc-63, acr-8, ric-3.1, unc-50 and unc-74). Parts of this work contributed to the PhD theses of TBD and JDN.

## Supplementary Material

Supplemental Figure 1. Subunit concatemer design

Supplemental Figure 2. Models of subunit concatemers

Supplemental Table 1. Summary of current responses

## Data Availability Statement

The data that supports this paper is available from R. Beech upon request.

---

**Supplemental Figure 1.**
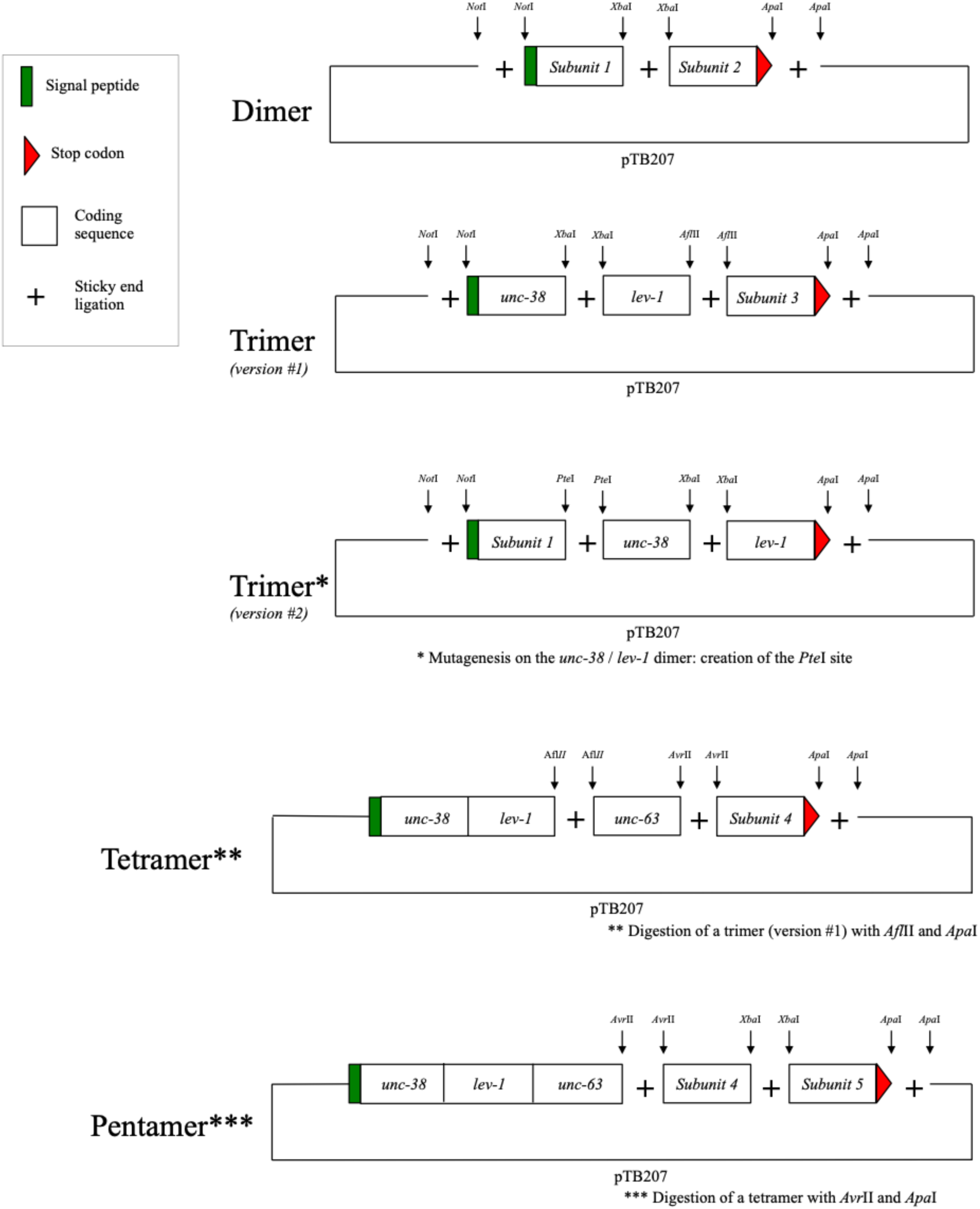
Subunit concatemer design. Subunit concatemers were made into one continuous construct. The C-terminal tail of the first subunit was linked to the N-terminal domain of the next subunit. Only the first subunits’ signal peptide and the last subunits’ stop codon were retained, making a single open reading frame and restricting the subunits to one specific configuration. UNC-63 required the addition of five Ser-Ala-Thr repeats, chosen as small, non-branched, hydrophylic residues added to its C-terminal tail.

**Supplemental Figure 2.**
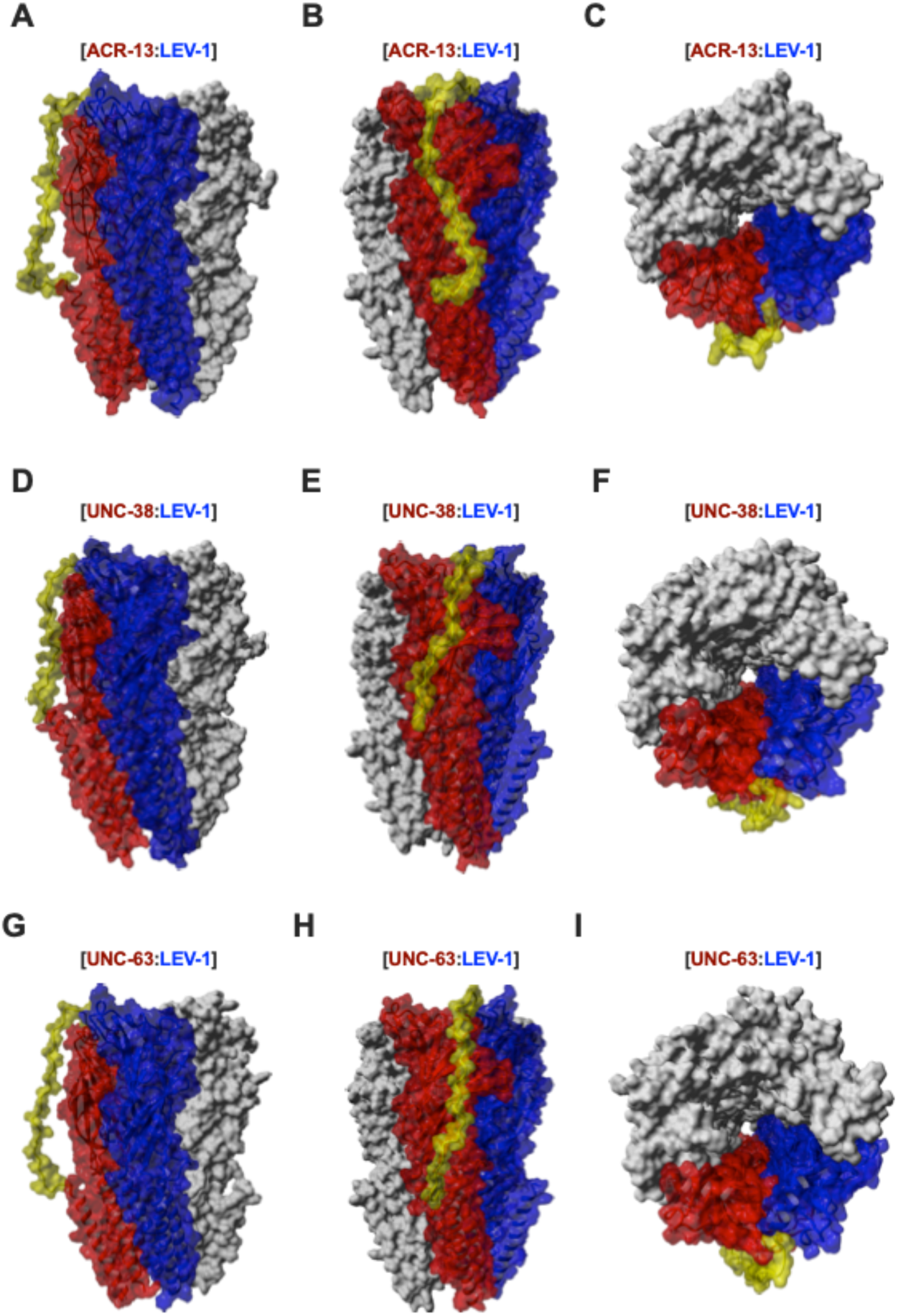
Model of subunit concatemers. Structural models of subunit dimers: (A – C) ACR-13:LEV-1; (D - F) UNC-38:LEV-1; and (G - I) UNC63-:LEV-1. Models were based on subunits A and B of the human alpha4/beta2 acetylcholine receptor (PDB: 5KXI) (Morales-Perez et al., 2016). The alignment was constructed by hand, based on the conserved secondary and tertiary structural elements. Red is the amino end subunit of the dimer, blue is the carboxy end subunit of the dimer and the amino acids between the end of TM4 on the red subunit and beginning of the blue subunit are colored in yellow (ie the linked region). UNC-63 is the only one where an artificial linker was added in. The models were developed using the hm_build macro of YASARA 23.12.24 (Krieger et al., 2002). In each case, the C-terminal of the principal face subunit was covalently attached to the N-terminal of the complementary subunit face.

**Supplemental Table 1.** Summary of current responses. Receptor subunit combinations are shown in response to 100 μM ACh and 100 μM LEV. C corresponds to *C. elegans* and H corresponds to *H. contortus*. Subunit numbers indicate the gene name number (example 38 = *unc-38*, etc). Error represents standard error. n.s: non-significantly different according to an unpaired Student’s t-test with a P-value p<0.05. * p< 0.05; **p<0.005; ***p<0.0002, **** p<0.0001. ^a^ values obtained from Duguet et al., 2016.

| Condition | ACh Current (nA) | n | LEV Current (nA) | n |
| --- | --- | --- | --- | --- |
| Cel-L-AChR-1 (with ACR-13) <sup>a</sup> | 1017 ± 128.6 | 10 | 518.8 ± 118.9 | 6 |
| Cel-L-AChR-2.1 (with ACR-8 & UNC-29) | 387.6 ± 34.37 | 10 | 80.90 ± 10.97 | 10 |
| Hco-L-AChR-1.2 (with UNC-29.2) <sup>a</sup> | 0.00 ± 0.00 | 9 | 0.00 ± 0.00 | 9 |
| [C38:C1] + C63 + C13 + C29 | 1149 ± 121.3<br>n.s.; p = 0.4682 | 9 | 618.3 ± 84.72<br>n.s., p = 0.4954 | 9 |
| [C1:C63] + C38 + C13 + C29 | 404.4 ± 88.31<br>*** | 12 | 199.6 ± 45.13<br>** | 12 |
| [C38:C1:C63] + C13 + C29 | 460.5 ± 81.40<br>*** | 20 | 204.2 ± 32.02<br>** | 17 |
| [C38:C1:C63:C13] + C29 | 36.96 ± 6.65<br>*** | 4 | 37.28 ± 2.46<br>* | 4 |
| C38 + C1 + C63:linker + C13 + C29 | 689.9 ± 221.6<br>n.s.; p = 0.1940 | 5 | 372.9 ± 189.1<br>n.s.; p = 0.5156 | 5 |
| [C63:C29] + C38 + C13 + C1 | 0.00 ± 0.00<br>**** | 8 | 0.00 ± 0.00<br>*** | 8 |
| [C63:C1] + C38 + C13 + C29 | 0.00 ± 0.00<br>**** | 8 | 0.00 ± 0.00<br>*** | 8 |
| [C38:C29] + C63 + C13 + C1 | 0.00 ± 0.00<br>**** | 8 | 0.00 ± 0.00<br>*** | 8 |
| [C13:C29] + C38 + C63 + C1 | 0.00 ± 0.00<br>**** | 8 | 0.00 ± 0.00<br>*** | 8 |
| [C13:C1] + C63 + C38 + C29 | 0.00 ± 0.00<br>**** | 8 | 0.00 ± 0.00<br>*** | 8 |
| [C63:C38] + C1 + C13 + C29 | 0.00 ± 0.00<br>**** | 8 | 0.00 ± 0.00<br>*** | 8 |
| [C63:C13] + C1 + C38 + C29 | 0.00 ± 0.00<br>**** | 8 | 0.00 ± 0.00<br>*** | 8 |
| [C38:C63] + C1 + C13 + C29 | 0.00 ± 0.00<br>**** | 8 | 0.00 ± 0.00<br>*** | 8 |
| [C38:C13] + C1 + C63 + C29 | 0.00 ± 0.00<br>**** | 8 | 0.00 ± 0.00<br>*** | 8 |
| [C13:C63] + C1 + C38 + C29 | 0.00 ± 0.00<br>**** | 8 | 0.00 ± 0.00<br>*** | 8 |
| [C13:C38] + C1 + C63 + C29 | 0.00 ± 0.00<br>**** | 8 | 0.00 ± 0.00<br>*** | 8 |
| [C38:C1:C13] + C63 + C29 | 0.00 ± 0.00<br>**** | 8 | 0.00 ± 0.00<br>*** | 8 |
| [C63:C38:C1] + C13 + C29 | 0.00 ± 0.00<br>**** | 8 | 0.00 ± 0.00<br>*** | 8 |
| [C13:C38:C1] + C63 + C29 | 0.00 ± 0.00<br>**** | 8 | 0.00 ± 0.00<br>*** | 8 |
| [C29:C38:C1] + C13 + C63 | 0.00 ± 0.00<br>**** | 8 | 0.00 ± 0.00<br>*** | 8 |
| [C38:C1:C63:C29] + C13 | 0.00 ± 0.00<br>**** | 5 | 0.00 ± 0.00<br>**** | 5 |
| [C38:C1:C63:C13:C29] | 0.00 ± 0.00<br>**** | 8 | 0.00 ± 0.00<br>*** | 8 |
| [C38:C1:C63:C29:C13] | 0.00 ± 0.00<br>**** | 8 | 0.00 ± 0.00<br>*** | 8 |
| [C38:C1] + C63 + C29 + C13 | 209.9 ± 37.42 | 9 | 83.23 ± 18.68 | 9 |
| [C38:C1] + C63 + C29 + C8 | 361.9 ± 73.63 | 10 | 71.15 ± 15.14 | 10 |
| [C38:C29] + C63 + C29 + C13 | 14.59 ± 2.06<br>*** | 7 | 7.51 ± 0.90<br>** | 7 |
| [C38:C29] + C63 + C29 + C8 | 66.90 ± 12.38<br>** | 8 | 16.08 ± 2.12<br>** | 8 |
| [C38:C29] + C63 + C1 + C13 | 0.0 ± 0.00<br>**** | 10 | 0.0 ± 0.00<br>*** | 10 |
| [C38:C29] + C63 + C1 + C8 | 0.0 ± 0.00<br>*** | 8 | 0.0 ± 0.00<br>*** | 8 |
| [C38:C29] + C63 + C13 | 0.0 ± 0.00<br>**** | 10 | 0.0 ± 0.00<br>*** | 10 |
| [C38:C29] + C63 + C8 | 0.0 ± 0.00<br>*** | 8 | 0.0 ± 0.00<br>*** | 8 |
| C38 + C1 + C63 + C29 + H8 | 955.5 ± 108.4<br>n.s.; p=0.7232 | 9 | 1008.8 ± 113.3<br>* | 9 |
| C38 + C63 + C29 + H8 | 383.9 ± 51.17<br>*** | 10 | 420.7 ± 62.23<br>n.s.; p=0.4345 | 13 |
| C38 + C1 + C63 + H8 | 0.0 ± 0.00<br>**** | 13 | 0.0 ± 0.00<br>**** | 13 |
| C29 + H38 + H63 + H8 | 527.2 ± 157.9<br>* | 7 | 369.1 ± 49.83<br>n.s.; p = 0.3064 | 5 |
| C1 + H38 + H63 + H8 | 119.4 ± 25.59<br>**** | 8 | 19.82 ± 2.51<br>*** | 8 |
| C13 + H38 + H63 + H29.1 | 2179 ± 200.1<br>*** | 12 | 1875 ± 213.3<br>**** | 11 |
| C38 + C1 + C63 + C13 + H29.1 <sup>a</sup> | 471.1 ± 51.48<br>** | 9 | 279.9 ± 34.02<br>* | 11 |
| C38 + C1 + C63 + C13 + H29.2 <sup>a</sup> | 28.61 ± 5.17<br>**** | 10 | 16.87 ± 3.83<br>**** | 10 |
| C38 + C1 + C63 + C13 + H29.3 <sup>a</sup> | 109.6 ± 20.88<br>**** | 9 | 38.93 ± 7.03<br>**** | 12 |
| C38 + C1 + C63 + C13 + H29.4 <sup>a</sup> | 49.20 ± 14.50<br>*** | 5 | 14.63 ± 2.64<br>** | 6 |
| C38 + C63 + C13 + H29.1 | 1766.1 ± 85.6<br>*** | 10 | 1637.3 ± 103.9<br>**** | 10 |
| C38 + C63 + C13 + H29.2 | 0.0 ± 0.0<br>**** | 8 | 0.0 ± 0.0<br>*** | 8 |
| C38 + C63 + C13 + H29.3 | 337.4 ± 72.86<br>*** | 9 | 73.24 ± 15.51<br>**** | 14 |
| C38 + C63 + C13 + H29.4 | 346.9 ± 84.10<br>** | 6 | 101.3 ± 31.80<br>*** | 11 |
| H38 + H29.2 + H63 + H8 + C1 | 236.0 ± 23.89<br>**** | 12 | 227.9 ± 17.76<br>** | 12 |
| H38 + H29.2 + H63 + C13 | 0.00 ± 0.00<br>**** | 8 | 0.00 ± 0.00<br>*** | 8 |
| H38 + H29.2 + H8 + C63 | 0.00 ± 0.00<br>**** | 10 | 0.00 ± 0.00<br>**** | 10 |
| H29.2 + H63 + H8 + C38 | 0.00 ± 0.00<br>*** | 5 | 0.00 ± 0.00<br>** | 5 |
| H29.2 + H8 + C38 + C63 | 0.00 ± 0.00<br>**** | 8 | 0.00 ± 0.00<br>*** | 8 |
| H29.2 + H8 + C38 + C63 + C1 | 156.1 ± 39.80 | 7 | 169.9 ± 27.07 | 6 |
|  | **** |  | * |  |
| Chim-1 + C13 + C38 + C63+ C1 | 23.07 ± 3.08 | 9 | 11.94 ± 1.78 | 9 |
| Chim-1 + C13 + C38 + C63 | 0.00 ± 0.00<br>**** | 7 | 0.00 ± 0.00<br>**** | 7 |
| Chim-1 + H38 + H8 + C63 | 0.00 ± 0.00<br>**** | 11 | 0.00 ± 0.00<br>**** | 11 |
| Chim-2 + C13 + C38 + C63 + C1 | 245.5 ± 37.84 | 19 | 173.2 ± 32.58 | 19 |
| Chim-2 + C13 + C38 + C63 | 16.27 ± 2.32<br>*** | 9 | 11.74 ± 1.9<br>** | 9 |
| Chim-2 + H38 + H8 + C63 | 39.87 ± 17.37<br>** | 6 | 53.33 ± 17.75<br>p = 0.0561 | 6 |

